# Unveiling Clonal Cell Fate and Differentiation Dynamics: A Hybrid NeuralODE-Gillespie Approach

**DOI:** 10.1101/2024.07.08.602444

**Authors:** Mingze Gao, Melania Barile, Shirom Chabra, Myriam Haltalli, Emily F. Calderbank, Yiming Chao, Elisa Laurenti, Bertie Gottgens, Yuanhua Huang

**Author notes:** These authors contributed equally (Mingze Gao, Melania Barile).

## Abstract

Recent lineage tracing single-cell techniques (LT-scSeq), e.g., the Lineage And RNA RecoverY (LARRY) barcoding system, have enabled clonally resolved interpretation of differentiation trajectories. However, the heterogeneity of clone-specific kinetics remains understudied, both quantitatively and in terms of interpretability, thus limiting the power of barcoding systems to unravel how heterogeneous stem cell clones drive overall cell population dynamics. Here, we present CLADES, a NeuralODE-based framework to faithfully estimate clone-specific kinetics of cell states from newly generated and publicly available human cord blood LARRY LT-scSeq data. By incorporating a stochastic simulation algorithm (SSA) and differential expression gene (DEGs) analysis, CLADES yields cell division dynamics across differentiation timecourses and fate bias predictions for the early progenitor cells. Moreover, clone-level quantitative behaviours can be grouped into characteristic types by pooling individual clones into meta-clones. By benchmarking with CoSpar, we found that CLADES improves fate bias prediction accuracy at the meta-clone level. In conclusion, we report a broadly applicable approach to robustly quantify differentiation kinetics using meta-clones while providing valuable insights into the fate bias of cellular populations for any organ system maintained by a pool of heterogeneous stem and progenitor cells.

## Introduction

One of the key challenges in developmental biology is to understand the complex cellular dynamics as well as the temporal ordering of cell states^1,2^. The interplay between cell proliferation (cellular expansion) and differentiation (phenotypic transition) plays a crucial role in various biological processes, including tissue development, regeneration, and the activation of innate immune response mechanisms^3^.

In recent years, the transcriptome-wide single-cell RNA sequencing (scRNA-seq) technique has emerged as a scalable approach for studying cellular trajectories, either on a snapshot of a transitioning cell population or via a time-series design. Computational strategies for dynamical analyses include both pseudotime-based trajectory inference^4–8^ and unspliced RNA-based RNA velocity methods for predicting gene expression profiles and their temporal changes^9–11^. However, scRNA-seq alone cannot provide fine-grained insights into clonal heterogeneity within cell clusters. Emerging lineage tracing techniques utilize unique and heritable DNA barcodes to track individual cells, offering a complementary approach to studying cellular dynamics^1,3,12^.

Techniques of lineage-tracing coupled with single cell sequencing (LT-scSeq) include retrospective analyses via endogenous genetic barcodes (e.g., mitochondrial variants and copy number variations^13–16^) and prospective designs with recently developed barcoding technologies via exogenous barcodes. Prospective designs can be broadly categorized into static barcoding with a one-off introduction to focus more on clone-specific differentiation^17,18^, and dynamical barcoding with multiple-time introductions to trace phylogenies^19,20^. In this study, we primarily focused on modeling static barcoding LT-scSeq data, where numerous (often heterogeneous) progenitor cells are labeled at an early point with a unique barcode that is then propagated to all other cell populations (progeny from here on), hence facilitating the delineation of a highresolution differentiation topology. One prominent example of this type of LT-scSeq technologies is the lentivirus-based system^21,22^ (also known as LARRY^18^), which has been recently employed, for example, for predicting clonal fate differentiation bias in hematopoiesis^18^ and mouse brain formation^23,24^, unveiling novel regulators/markers involved in reprogramming^25^ and cell differentiation^26^, and identifying pathways relevant to cancer progression^27,28^.

Several computational algorithms have been proposed to facilitate the analysis of population dynamics and LT-scSeq experiments recently. Depending on the space of the cell states, one may group these methods in different categories. First, for continuous model, Fischer et al.^29^ developed pseudo-dynamics, which could model population distribution shifts to quantify developmental potentials for time-series data. Second, a common choice is to mimic a near-continuous space by employing a finite state mapping where each cell is treated as a state. For instance, LineageOT^30^ maps cells from the same clone and two adjacent time points using optimal transport; it was developed to recover lineage couplings in CRISPR-based lineage tracing datasets and outperforms the original OT method that operates without barcodes^31^. Another example is CoSpar^32^, which performs topology mapping under the constraints of sparsity and coherence within clones. This method brought new insights to multi-clonal time series data analysis with respect to the identification of early cell fate bias at the cellular level. Third, discrete state space over cell types is also commonly used to ensure better interpretability, higher robustness, and computational efficiency^33^. Finally, other approaches have also significantly contributed to this field by modeling individual genes or phenotypes as a different type of cell state space, for example, upon learning smoothed transcription and regulatory dynamics^34^, or explicitly analyzing the potency bias during HSCs’ reactivation following platelet depletion^35^.

Despite various efforts to unveil more information from LT-scSeq data, several technical challenges persist, limiting the practical applicability of analysis tools. In addition to common issues such as the loss of barcodes or small clone size, scRNA-seq is destructive and only captures a fraction of the total cells in a dish, resulting in the possibility of tracking only the proportional changes of cell state abundance^32^ rather than the actual kinetic rates that affect the overall population dynamics. This hampers the exploration of clone-specific dynamical patterns. On the other hand, having reliable kinetic rates enables the reconstruction of differentiation topologies and provides a quantitative estimation of division dynamics^35^ as well as fate bias of progenitor cells, which is useful under various scenarios, e.g., investigation of stages of development or abnormal patterns related to aging and diseases.

Inspired by recent research efforts and to address the aforementioned problems, we developed a robust and generalizable computational algorithm to analyze LT-scSeq datasets focusing on a static barcoding system (LARRY, Fig. 1a). Our tool, CLADES (**C**lonal **L**ineage **A**nalysis with **D**ifferential **E**quations and **S**tochastic Simulations), comprises two key components: 1) *a model estimator*, namely a NeuralODE^36^ based framework, to delineate meta-clone specific trajectories and state-dependent transition rates; 2) *a data generator* via the Gillespie algorithm^37^, that allows a cell, for a randomly extracted time interval, to choose either a proliferation, differentiation, or apoptosis process in a stochastic manner. We estimated the summary of the divisions between progenitors and progeny, and showed that the fate bias between all progenitor-fate pairs can be inferred probabilistically. In conclusion, our work elucidates clone-specific dynamics and provides a quantitative description of the differentiation trajectory between a progenitor and its progeny.

**Figure 1:**
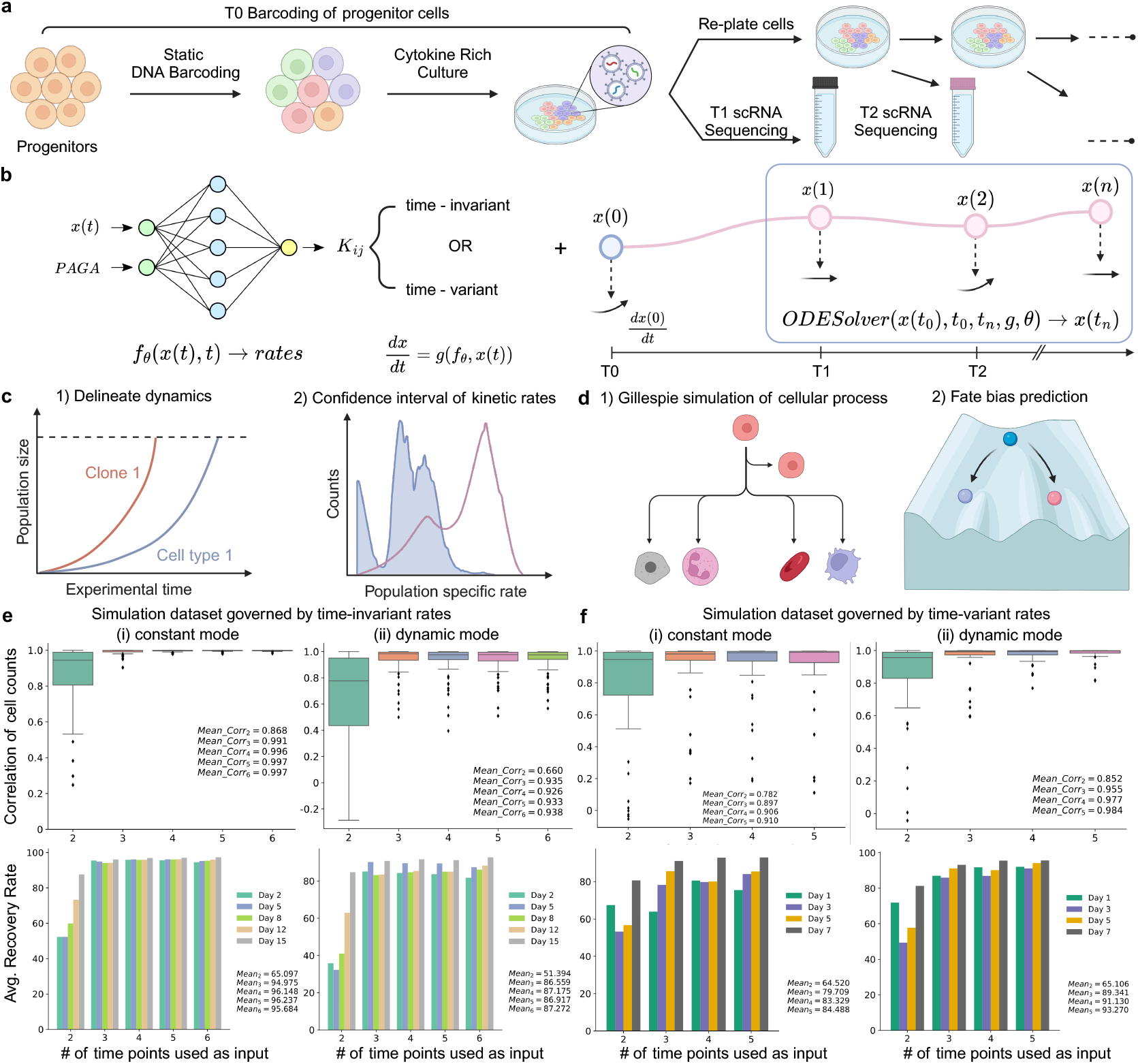
Overview of the experimental design of LT-scSeq, CLADES’s architecture and its robustness on synthetic datasets. **a**, General workflow of LT-scSeq experiment using static barcoding techniques with viral integration. DNA barcodes are induced at an early time point, whilst scRNA-seq data and clonal information are acquired at subsequent time points. **b**, CLADES takes total cell counts and transition directions as input, then uses MLP for estimating the transition rates between populations, and NeuralODE for reconstructing cell counts at each time point. **c**, CLADES is able to infer the dynamic changes of population size on various resolutions and output the associated CI of transition rates. **d**, With the estimated kinetic rates of each clone, CLADES can further: 1) simulate detailed topologies of division and differentiation, and 2) identify putative regulators/markers of cell fate bias of progenitor cells. **e, f** CLADES was validated in both constant mode and dynamic mode, that is, given the cell counts generated by either time-invariant or time-variant rates, the performance of both modes using only training time points as input was evaluated. **e**, Given cell counts generated by time-invariant rates, the constant mode (i) consistently performs better than its counterpart (ii) both on the correlation of cell counts (upper row) and on the average recovery rate (lower row) no matter how many time points were used by the model; vice versa, the dynamic mode performs better for cell counts generated by time-variant rates **f**, (i,ii). Upper row: correlation of cell counts on populations between generated and predicted values for each trial, where `Mean`_`corr`_`n` is the mean correlation for all the trials tested on *n* training time points. Lower row: average recovery rate in percentage view (see helping functions of Methods section) for each trial, where `Mean`_`n` is the mean correlation for all the trials tested on *n* training time points.

## Results

### ODE functions for clone-specific dynamics

We model clone dynamics as a system of independent ordinary differential equations (ODEs) to infer the time-specific transition rates between cell states. The dynamics of each clone (or each meta clone, namely a group of clones with similar dynamical profiles; see below) are described by the same equation but clone-specific parameters; this aligns with the assumption that only intra-clone transitions are allowed (cells keep the same clone identity throughout the whole experimental time). Therefore, the estimation and inference processes are independent for each clone. Without loss of generality, we described the (multivariate) ODE function *f* for a specific clone *c* with parameters *θ* as shown in Eq.(1),

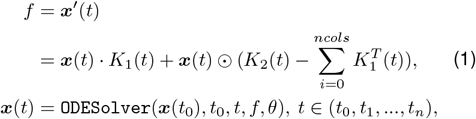

where · is the dot product of two matrices, ⊙ is elementwise multiplication, ***x***(*t*) is a vector of total counts for all cell states (interchangeably as cell populations, e.g., 12 cell types), and *t* refers to the real time of the biological system. *K*_1_(*t*) mimics the differentiation among populations and is based on the edges of the PAGA^38^ graph (denoted as *L*, Methods) with expert curation; it is a non-negative and strictly upper triangular matrix (Supplementary Fig. S1). *K*_2_(*t*) (diagonal of the transition rate matrix *L*) is a one-dimensional vector, representing the overall net proliferation rate (proliferation minus apoptosis) for each cell state. *K*_1,2_(*t*) can either be constant, which gives the classic population growth model with exponential change of population sizes, or a function of the real time *t* (Fig. 1b), which allows for more flexibility in our model^39–41^.

The model is designed for time series LT-scSeq experiments (Fig. 1a-b) and requires three types of input data: 1) the estimated total cell counts ***x***(*t*) per time point *t*_*i*_, clone *c*_*j*_ and population *p*_*k*_; 2) putative transition directions between populations, usually derived from the PAGA graph with expert curation, which incorporates prior knowledge into the model; 3) a binary vector indicating terminal/progenitor cell populations. Under the assumption that cellular divisions and differentiations between distinctive states are a stochastic process governed by a set of transition rates, we then interpolate the total cell counts on intermediate time points using NeuralODEs with biologically informed constraints (Fig. 1b, Methods). This involves solving an optimization problem between observed data and model predictions with meaningful penalties.

CLADES takes the input data and the putative terminal cell states as prior information, then feeds them into a Multi-Layer Perceptron (MLP, Methods) with 2 layers, from which the model outputs the rate transition matrices (Supplementary Note 1) among populations and the predicted cell counts using an ODE solver. The rates can be timeinvariant or time-variant depending on the user’s choice. After calculating the rates, CLADES can reconstruct the dynamic changes of cell counts at both the population and the clone levels and derive the associated confidence intervals (CI) of the kinetic transition rates (Fig. 1c) as well, providing a measure of uncertainty. By means of the Gillespie algorithm (Supplementary Note 2, Methods), CLADES also provides a quantitative summary of cell divisions, assesses the potential fate bias of progenitors, and explores the link with the DEGs (Fig. 1d).

### Robustness of CLADES on synthetic datasets

In order to evaluate the performance of CLADES in both constant and dynamic modes, we generated synthetic datasets with time-invariant and time-variant transition rates.

First, we applied CLADES on a time-invariant synthetic dataset (given by constant rate ODE functions), where the time-invariant rates for each clone serve as the ground truth and cell counts are governed by Eq.(1). To test whether CLADES can recover the correct dynamics and provide guidance for appropriate usage with LT-scSeq experiments, we conducted 5 independent trials with different sampling intervals of synthetic data as training sets and used 5 unobserved time points as testing sets to prevent data leakage, for both constant and dynamic modes of the model (Supplementary Table S1).

As expected, the performance of CLADES improved as the available time points increased, eventually reaching a plateau at around 3∼4 time points in this time-invariant setting (for both constant and dynamic modes; Fig. 1e). Since the total cell counts are dependent on the transition rates, we observed a similar trend in inferred time-invariant rates on the testing set as well (Supplementary Fig. S2a). Also, we found that generally, the constant mode performs better than the dynamic mode because the synthetic data was synthesized based on it, indicating the non-negligible risk of over-fitting for the dynamic mode when the pattern of data is extremely clean and follows an ideal time-invariant rate scenario (Fig. 1e).

Second, with the time-variant scenario, we demonstrated the limitations of the constant mode when facing complex data structures which are prevalent in most realworld scenarios, and highlighted the flexibility and robustness of the dynamic mode. Specifically, similar synthetic settings were adopted (Supplementary Table S2), whilst this time the data was generated using time-variant rates. For both the constant and dynamic modes of CLADES, the accuracy increased when more training time points were given and an obvious plateau had not been observed (Fig. 1f). However, the performance in inferring the correct rates is not prominent for the constant mode when the size of the training dataset increases (Supplementary Fig. S2b). Furthermore, considering as empirically good a Pearson correlation *r >* 0.95, we found that the constant mode is heavily under-fitted even when more training data was provided and it could not recapitulate the correct dynamics. Interestingly, we also noticed that, apart from outperforming the constant mode when applied to its paired data (Fig. 1e (i), Fig. 1f (ii)), the dynamic mode generally yields better results even when it is applied to data generated via the other mode (Fig. 1e (ii), Fig. 1f (i)), which suggests that the dynamic mode is more robust to the “wrong” model and therefore could be a good default choice. As for the experimental design, we recommend using at least 3 time points to secure the desired results.

In conclusion, for datasets with simpler patterns, the constant mode is sufficient with tolerable errors, whilst for other biological systems that are more sophisticated, the dynamic mode of CLADES is preferred.

### Characterizing kinetic rate differences in human cord blood hematopoiesis

We applied CLADES to a newly generated in vitro LARRY-based LT-scSeq data with 3 time points (68,856 high-quality cells with 24,885 being barcoded; see details in Supplementary Table S3 and Supplementary Note 3). DNA barcodes were introduced at Day 0 in cord blood CD34+ cells and sampled for sequencing at Day 3, 10, and 17 (Fig. 2a, Fig. 2b right). Upon incubation, a group of initial hematopoietic stem and progenitor cells (HSPCs) differentiated into 12 distinct cell populations of progenitor or mature blood cells (i.e., cell types; Fig. 2b left). Here we define clone as a group of cells carrying the same barcode. Because a significant portion of the barcodes collected (3,334 out of 3,940) were low-quality clones (i.e., containing one or two cells along the entire time course), they were filtered out during the preprocessing step (Fig. 2d, Methods).

**Figure 2:**
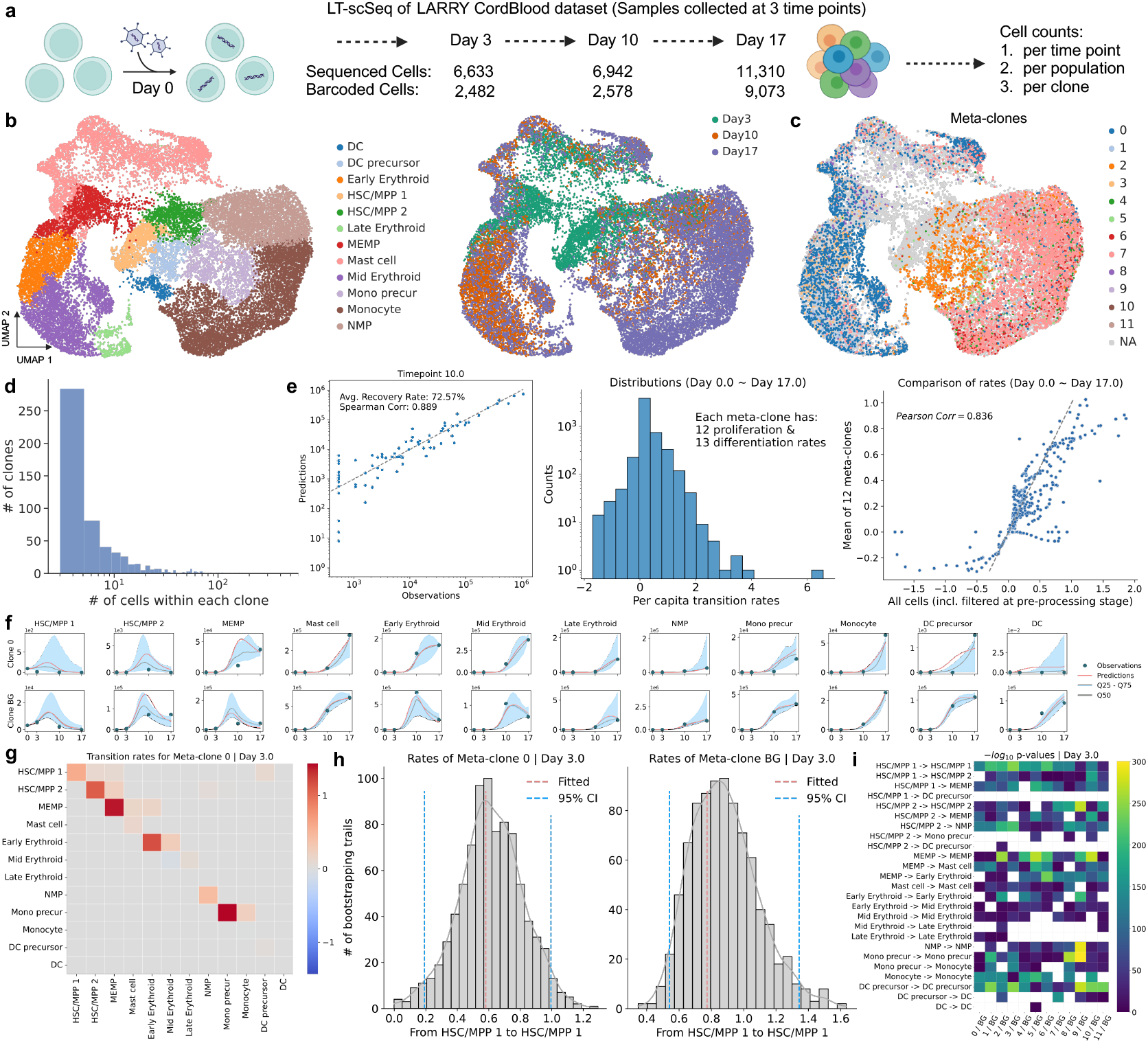
CLADES accurately reconstructed the clone-specific dynamics on a LARRY LT-scSeq data of human cord blood development system. **a**, Schematic of the data collection procedure for our newly generated LARRY human cord blood data. The model takes total cell counts as inputs. **b**, UMAP illustration of the dataset, which contains 12 populations and 3 sequenced time points. **c**, Combined UMAP for the distribution of each meta-clone on the same landscape. NA represents non-barcoded cells. **d**, Histogram of the number of cells belonging to each barcode shows the sparsity and heterogeneity of clones, making the estimation of clone-specific dynamics infeasible and leading to the aggregation of individual clones into meta-clones. **e**, Left: CLADES accurately recovered the real cell counts at different time points. Middle: Proliferation and differentiation rates estimated by CLADES are within a reasonable range. Right: the mean transition rates of all meta-clones given by CLADES resemble rates of the overall biological system. **f**, Example of observed and fitted cell counts and estimation error for meta-clone 0 and background cells using the dynamic mode of CLADES. The red line is the originally fitted curve whilst the light blue region represents the 25%−75% quantile. **g**, Transition rate matrix between populations of meta-clone 0 at Day 3, which is the graphical illustration of *K*_*ij*_ in ODE functions, illustrating that CLADES has the ability to infer rates at any given time point. **h**, 95% confidence interval of estimated transition rates between cell populations are given by the bootstrapping approach. Blue dashed lines are CI whilst the red dashed line is the original fitted value. **h** is an example extracted from **g** and **i. i**, Inferred rates given by bootstrapping were compared with each other using *p* values from the student t-test or Mann-Whitney U rank test and mean absolute change of parameters were used to find the most active (distinct) meta-clone for all transition pairs. Blank region means no significance or the corresponding population has not been produced at all.

DNA barcodes with few cell counts make the analysis of individual clones infeasible, therefore, to further reduce stochasticity and model complexity, clones were clustered into meta-clones (Supplementary Fig. S3) based on the similarity among their time and state-dependent number of barcoded cells (Fig. 2c, Supplementary Fig. S4). The assumption is that hematopoiesis can be conceptualised by a finite number of differentiation behaviours; therefore, we defined meta-clones as groups of clones with similar kinetic rates and cell counts. Notably, meta-clones demonstrate distinct preferences for terminal fates (e.g., Mast cell, Late Erythroid, Monocyte and DC) decisions, indicating strong fate bias among HSC/MPP 1 population (Fig. 2c, Supplementary Fig. S5, Table. S5). For instance, meta-clone 7 predominantly biased towards the Monocyte lineage, whilst meta-clone 0 had a preference for the Erythroid lineage. The number of meta-clones is a hyperparameter that can be adjusted to explore the data at different levels of resolution.

Given expert-curated putative transition directions derived from PAGA (Supplementary Fig. S1), CLADES successfully reconstructed the total cell counts at the experimental time points (Fig. 2e left, Supplementary Fig. S6a, b) and interpolated counts on unknown time points along the entire trajectory using both constant and dynamic modes while also providing associated estimation errors (Fig. 2f, Supplementary Fig. S7). Notably, the dynamic mode proved to have better performance provided that proper constraints are enforced to prevent model overfitting (Supplementary Table S4).

As stated above (Eq. 1), the population balance between cell states is governed by transition rates which are the per capita change within a unit of time. As a result, the majority of clone-associated rates should lie within a feasible range (e.g., fit biological prior knowledge, Methods, Fig. 2e, middle). Moreover, the average behavior of all meta-clones should resemble the overall dynamics of the system, including both the barcoded and non-barcoded cells (background from here on, or BG in short), Fig. 2e, right. The output of CLADES includes a transition rate matrix among cell states for any meta-clone and time stamp (Fig. 2g). It provides insights into the heterogeneity of transition rates among meta-clones and offers partial evidence of the distinct behaviors of meta-clones.

The parameters returned by CLADES are point estimates that lack uncertainty. We then used bootstrapping (Methods) to estimate the 95% confidence intervals (CI) of the transition rates (Fig. 2h). We used statistical tests (the student t-test or Mann-Whitney U rank test) to assess the statistical significance of dynamical differences among meta-clones. It is worth noting that, when performing the tests, we took precautions to avoid abnormal significance in the bootstrapping results, especially for populations with zero cell counts. Therefore, we introduced the mean absolute change of parameters as an additional criterion (Methods, Fig. 2i shows comparison with background, extracted from Supplementary Fig. S8). From the perspective of developmental trajectory dynamics, distinct clones can exhibit differences in rates at specific stages of progenitors/precursors. At Day 3, for example, meta-clone 9 were found to have strong statistically different rates compared with the background in the Monocyte lineage (Fig. 2i).

### Resolving cell division history and fate decision bias of progenitor cells

After estimating the transition rates between cell states using CLADES, we then employed the Gillespie algorithm (1, a stochastic simulation algorithm originally used to depict chemical reaction processes) to delineate the behavior of the clones making up our system. A typical workflow of a Gillespie simulation starts with a single progenitor (HSC/MPP 1 for this dataset). For each step, the time interval until the next reaction (proliferation, differentiation, or apoptosis) is extracted from an exponential distribution (where the parameter *λ* is a staterate-dependent value); then, a reaction is picked to occur based on the currently available cell states and the previously estimated transition rate matrices. The simulation continues and updates cell states until certain stopping criteria are met (Fig. 3a).

**Figure 3:**
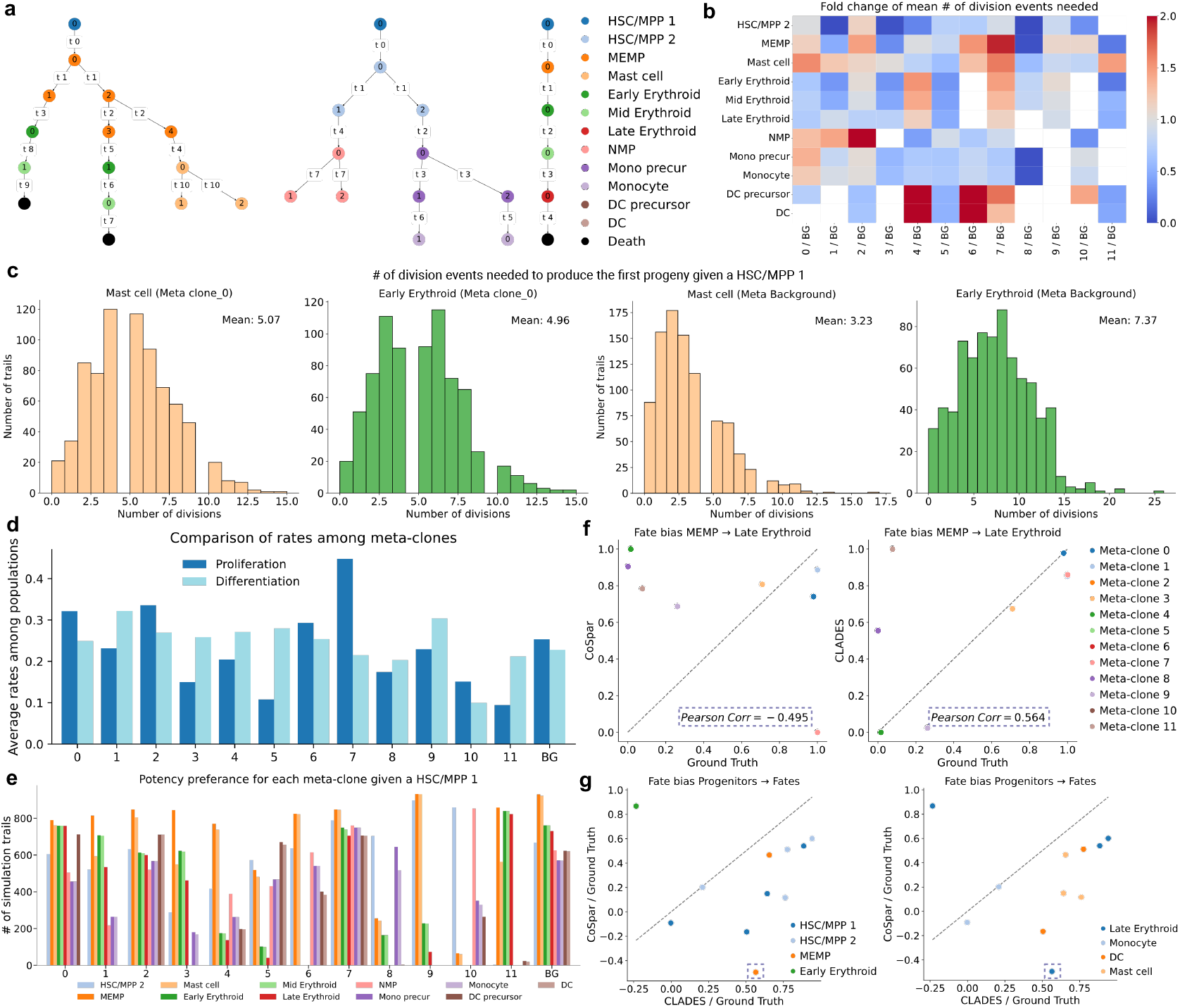
Gillespie algorithm reveals division summary of progenitor populations and more accurate fate decision bias. **a**, Examples of lineage trees resulting from the Gillespie algorithm; each simulation trail starts with one HSC/MPP 1 cell only. Nodes represent cell ID, that is, its order of generation. The edge label is the order of occurred reactions. **b**, Overall fold change of mean number of divisions needed to produce the first progeny starting from one HSC/MPP 1 for each meta-clone compared with background. **c**, Examples of the number of division events needed to produce the first Mast cell or the first Early Erythroid starting from one HSC/MPP 1 for meta-clone 0 (i) and background cells (ii) respectively. **d**, The average proliferation and differentiation rates for each meta-clone correspond to the abundance of total population size and potency potential of progenitor cells. **e**, Bar plot shows the potency preference and fate bias for each progeny of HSC/MPP 1 among all meta-clones. E.g., meta-clone 0, 2, and 7 are multi-potent clones whilst meta-clone 9, 10, and 11 are more likely to be uni-potent clones. The *y* axis indicates number of simulated trails a certain progeny has been generated out of all simulations (1,000 in our analysis). **f**, An example of fate decision bias comparison between CoSpar, CLADES, and GroundTruth for each meta-clone. The transition pair used for benchmarking is from MEMP to Late Erythroid. Note that for a fair comparison, the value is scaled using min-max normalization. Therefore, zero does not represent no bias towards that fate, instead, it means relatively smaller bias than other meta-clones. **g**, Combined benchmark results of all progenitor fate transition pairs for CLADES and CoSpar. Left panel, progenitors. Right panel, fates. The plot is formed based on the correlation scores of each transition pair (as shown in the dashed square of panel **f** as an example).

We ran Gillespie on all meta-clones and for each of them we inferred several properties, of which the most relevant are: 1) the number of cell division events that occurred between the original HSC/MPP 1 and the first cell produced for each progeny (Supplementary Fig. S9); 2) the number of trials where a certain population was produced out of all possible simulations trials (1,000 in this case); 3) the fate decision bias of each meta-clone towards the different terminal states. In other words, our simulations yielded a quantitative summary of differentiation topologies and terminal state preferences for each meta-clone. For instance, for meta-clone 0, it takes around 5 division events on average to produce the first Mast cell, significantly higher than that of background with 3 divisions, whilst HSC/MPP 1 differentiates into Early Erythroid faster than that of the background (5 vs 7 divisions; Fig. 3c). This indicates that meta-clone 0 has a faster route for generating erythroid lineage compared with the average behavior of all cells and vice versa for Mast cells. Supplementary Fig. S9 illustrates the comprehensive comparison of all other cell populations. Taken all the simulation results together (Supplementary Fig. S10a), we could compare the differences in fold change among all meta-clones or with respect to the background cells (Fig. 3b, Supplementary Fig. S10b).

Meta-clones have distinct differentiation preferences (Fig. 2c, Supplementary Fig. S5, Table. S5). Some are multi-potent, for instance, meta-clones 0, 2, 7 have 4 terminal states and meta-clones 1, 3 have 3, whilst others are either bi-potent or uni-potent, being committed to a specific lineage such as Mast cells, Late Erythroids, or Monocytes. We also noticed that a higher proliferation rate normally corresponds to meta-clones with larger population sizes; therefore meta-clones with a similar fate realisation, but different sizes, can also bear differential kinetic rates (Fig. 3d, Supplementary Table S5). Average rates are calculated using all available proliferation/differentiation rates spanning from Day 0 to Day 17. Overall, we saw that potency preferences are highly meta-clone specific, and that the bias towards each progeny (including both intermediate and terminal states) given a HSC/MPP 1 can be inferred probabilistically using Gillespie simulation (Fig. 3e).

Next, we were interested to see whether the fate bias of the progenitor cells can be predicted at the earliest experimentally collected stage (Supplementary Fig. S11a). CLADES was compared with CoSpar^32^, the state-of-the-art method to predict the fate bias of ancestor cells based on LT-scSeq data. However, since CLADES operates at the meta-clone scale instead of the scale of individual cells, for fair comparison, we pooled the fate bias values of each cell generated by CoSpar (Supplementary Fig. S11b, Methods) to the same meta-clone level for each progenitor. By using the number of unique barcodes in a terminal population and all cell states along the transition path as ground truth (Methods), CLADES generally outperformed CoSpar in predicting fate bias on the meta-clone level and had a higher Pearson correlation for all meta-clones (e.g., MEMP *→* Late Erythroid, Fig. 3f). We benchmarked CoSpar and CLADES on all possible progenitor-fate pairs by means of the Pearson correlation between the candidate algorithms and the ground truth (Fig. 3g). CLADES consistently showed improved performance on 9 out of 11 transition pairs. Of note, although CLADES shows higher accuracy in predicting cell fate at a meta-clone resolution, it is still not applicable at the individual cell level and interpolation approaches, e.g., via neighborhood graph, require further evaluation (see Discussions).

### CLADES recapitulates the cellular dynamics of murine hematopoiesis

We applied CLADES to a publicly available mouse hematopoietic dataset that was introduced by Weinreb, et al.^18^ and also used to benchmark CoSpar^32^. This dataset contains a larger number of cells compared with the cord blood dataset, allowing us to explore the gene signatures corresponding to differential rates. Compared with the cord blood data (4 terminal fates out of 12 cell states), there are now more possible potential fates (10 out of 22, Supplementary Table S8), while the number of time points is the same (3, Fig. 4a left and middle panel).

**Figure 4:**
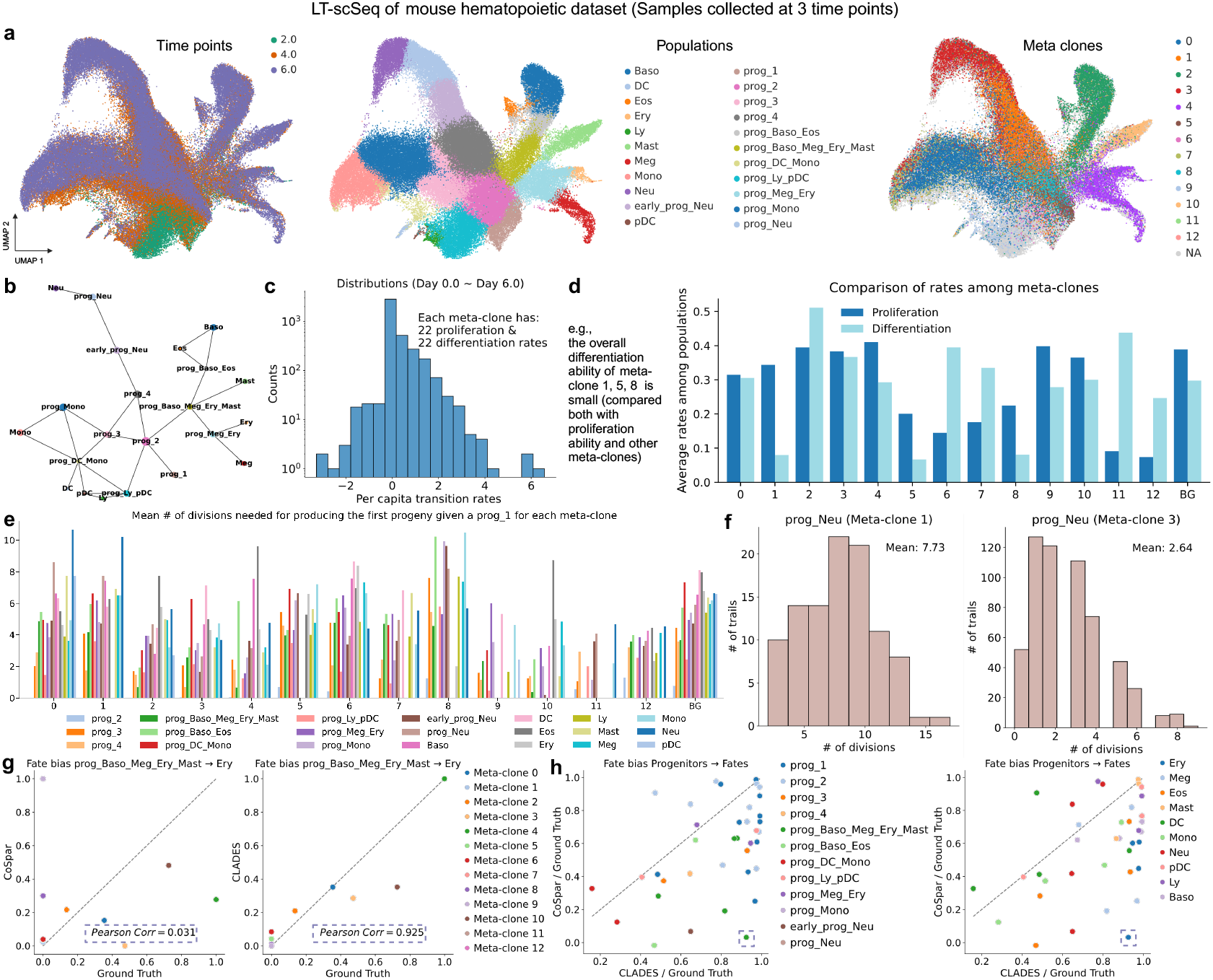
CLADES captures the key dynamics of mouse hematopoietic system and provides better fate bias predictions. **a**, UMAP illustration of the mouse hematopoietic dataset which contains 3 time points and 22 manually defined populations (Left, Middle). We defined 13 meta-clones based on time and state-dependent number of barcoded cells (Right). **b**, The putative transition directions were given by the PAGA graph with expert curation which acted as a guide for the model. **c**, The overview of the estimated proliferation and differentiation rates. **d**, Average rates are tightly associated with the population size of meta-clones, e.g., meta-clones 5 and 8 have a large gap between proliferation and differentiation rates, and they hardly produce differentiated cells. **e**, Mean number of divisions needed to produce the first progeny starting from one prog_1 cell were derived from the Gillespie algorithm. Higher bars indicate a slower process, which is a reflection of bias to terminal fates. **f**, An example extracted from **e**. Both meta-clone 1 and 3 are biased for neutrophils lineage whilst meta-clone 3 is more active compared with meta-clone 1. **g**, Example of the comparison between fate biases as calculated by CoSpar and CLADES for the transition pair prog_Baso_Meg_Ery_Mast and Ery. Meta-clone 4 has the highest bias towards the erythroid lineage compared to other meta-clones and CLADES has a higher Pearson correlation compared with the ground truth. **h**, The overall benchmarking results for all progenitor-fate pairs, indicating that CLADES improves the fate bias prediction with a large margin at the meta-clone level. Each dot in the scatter plot comprises the correlation scores between two methods with ground truth. The same dot represents progenitors in the left panel and fates in the right panel.

This dataset has 5,859 unique barcodes in total, with some of them being multi-potent whilst others being unipotent. Of note, only 1,989 clones appeared in the terminal states (Supplementary Fig. S11c). This indicates the low capture rate of barcodes and justifies the necessity to merge individual clones into meta-clones. We followed the same preprocessing pipeline as the cord blood dataset and constructed 13 meta-clones using again the time and state-dependent number of cells in each clone (Fig. 4a right panel, Supplementary Fig S12). Since this dataset does not contain FASC-based counting, we used an estimated fold expansion of haeamtopoietic stem and progenitors cells in culture as an alternative to the total cell counts and scaling factors (Supplementary Table S6, Methods). We applied both constant and dynamic modes of CLADES using the PAGA graph with expert curation as guided transition directions (Fig. 4b). Interestingly, we found that both modes performed similarly, suggesting that, in this dataset, the kinetics of the in vitro system are nearly constant (Supplementary Table S7).

The distribution of estimated proliferation as well as differentiation rates provided by CLADES overall falls within a reasonable range (Fig. 4c), with the average among populations and time points reflecting the characteristics of each meta-clone (Fig. 4d, Supplementary Table S8). For instance, meta-clones 1, 5, and 8 do not produce many terminal cells, and their differentiation rates are smaller than most of the other meta-clones. Meta-clones 5 and 8 have a smaller population size with the majority of cells remaining as progenitor cells during the 6-day culturing period, consistent with their smaller estimated proliferation rates than that of meta-clone 1. With the same Gillespie analysis, we could quantify the fate bias for each terminal state given a progenitor cell, and we obtained the division summary for each meta-clone (Fig. 4e, Supplementary Fig. S14). For instance, meta-clone 2 clearly has a fast route towards basophil cells (pink bar) and has the lowest value compared with other meta-clones. Both meta-clones 1 and 3 have a strong bias towards the neutrophil lineage (Supplementary Table S8), whilst the major difference is that, given the same period of incubation, meta-clone 3 seems to be more active than meta-clone 1. This is also reflected by the Gillespie analysis: the mean number of division events needed to produce the first prog_Neu given a prog_1 in meta-clone 1 is around 8, whilst for meta-clone 3 is 3 (Fig. 4f).

For the benchmarking of fate bias prediction results with CoSpar, we followed similar settings as used in cord blood analysis. CoSpar produces a set of scores for each cell given a terminal fate population (as shown in Supplementary Fig. S11d for Monocytes and Erythroids lineage) and we pooled the scores for cells within the same metaclone to get the average fate bias for that meta-clone. Taking the transition pair of prog_Baso_Meg_Ery_Mast and Ery (Erythroid) as an example (Fig. 4g), CLADES accurately recovered the relative fate bias for the metaclones with a much higher correlation score. Specifically, the meta-clone with the strongest bias towards the Ery lineage given by CLADES is meta-clone 4, which aligns with the ground truth (Supplementary Table S8), whilst CoSpar assigned the highest value to meta-clone 9, which does not produce erythroid-related cells at all during the culturing period. The results for all possible progenitor-fate pairs are shown in Fig. 4h, which consistently shows the improved performance of CLADES (30 out of 37) in inferring fate bias of meta-clones.

### Clonal kinetics and characteristics of population sizes can be inferred by early coordinated signatures

We showed earlier that the different output in terms of timescales and fate realization of early progenitors unveils differences in transition rates. We thus seek to connect such differences to possible heterogeneity in the transcriptome signature, arguing that the latter may coordinate the transition rates towards each lineage and terminal fate. We first considered cells belonging to the prog_1 population from the mouse hematopoietic dataset, as prog_1 is the earliest progenitor population (Fig. 5a, left panel) and calculated its DEGs across all meta-clones (Fig. 5a, right panel, Supplementary Fig. S15, S16). We found metaclone-specific characteristic genes that have differential expression levels and that are expressed in different areas of the dimensionally reduced landscape (e.g., some progenitors are scattered around whilst others are clustered together).

**Figure 5:**
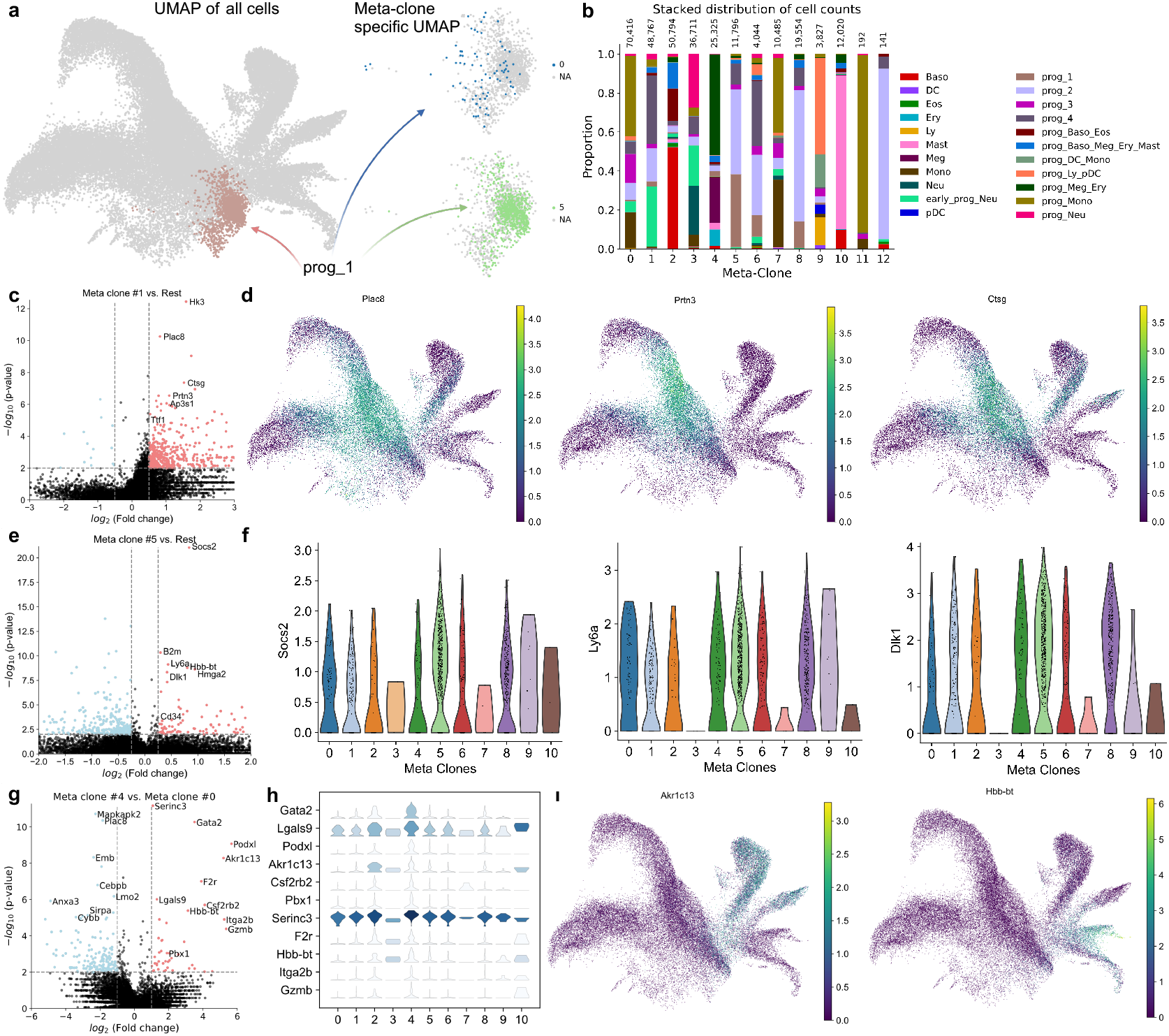
The fate bias and kinetics of each meta-clone is determined by its early transcriptomic signatures. **a**, Left, Only cells from the earliest progenitor populations were used for DEGs analysis, which is prog_1, as shown with brown color in the UMAP landscape for all cells. Right, examples of meta-clone-specific UMAPs. **b**, Stacked distribution of cell counts in percentage view where the *x* axis shows the meta-clones, whilst the *y* axis shows the proportion of cell counts for each population, and the number at the top of each bar is the total counts for each meta-clone. **c**, Volcano plot of DEGs between prog_1 cells within meta-clone 1 and all other prog_1 cells. Identified genes are either predominately over-expressed in the neutrophil maturation process (*Plac8, Prtn3, Cstg*) or affect its functions through associated pathways (*Hk3, Ap3s1, Ttf1*). **d**, Examples of gene expression UMAP plots on all cells for *Plac8, Prtn3* and *Cstg*, illustrating its over-expression in neutrophil lineage. **e**, Volcano plot of DEGs between prog_1 cells within meta-clone 5 and all other prog_1 cells. Identified genes are mostly associated with the stemness of a cell (e.g., *B2m, Hmga2*). **f**, Examples of gene expression violin plots on prog_1 cells for *Socs2, Ly6a* and *Dlk1*. The selected genes are generally expressed higher in meta-clone 5 than in other meta-clones. Interestingly, the existence of *Socs2* would inhibit the differentiation process, which explains why meta-clone 5 has such a low differentiation rate. **g**, Volcano plot of DEGs between prog_1 cells within meta-clone 4 and meta-clone 0. Both meta-clones are multi-potential whilst meta-clone 4 is more biased to Erythroid, Mast cell, Megakaryocyte, and Basal lineages compared with meta-clone 0 which is more biased to Monocytes. **h**, Staked violin plot for identified genes in **g. i**, Examples of UMAP expression profiles for *Akr1c13* and *Hbb-bt* on all cells.

Our hypothesis is that, even for the earliest progenitor cells, their preference for specific fate lineages is determined by subtle difference in transcriptomic profiles and could be accurately predicted. For instance, within prog_1, the DEGs of meta-clone 1 compared to all other metaclones are associated with neutrophil maturation (Fig. 5c), which is consistent with meta-clone 1 having a large population size in the neutrophil lineage (Fig. 5b). The upregulated genes either regulate the neutrophil maturation process or affect its functions through associated pathways. One example is *Plac8*, whose connections with neutrophils’ ability to kill phagocytosed bacteria have been reported^42,43^. Another identified DEG for this meta-clone is *Prtn3*, which confirms that this meta-clone is biased towards the Neutrophils lineage, because *Prtn3* belongs to NPSs (neutrophil serine proteases), a set of genes generally over-expressed in Neutrophils^44,45^ (Fig. 5d). Other identified genes like *Hk3*, might drive this meta-clone towards the neutrophil lineage because it is a key enzyme involved in the glucose phosphorylation process, a dominant metabolic pathway in neutrophils^46–48^. This suggests the predictable nature of terminal cell states within metaclones.

We also noticed that both meta-clones 5 and 8 are almost childless during the 6-day incubation period with extremely low differentiation rate to terminal states, with most of the cells staying in prog_1 and prog_2 stage and a few in prog_4 (Fig. 5b). To further analyze this unique behavior, we computed the DEGs (Supplementary Fig. S16) between meta-clone 5 and all other meta-clones within prog_1 cells (Fig. 5e). Interestingly, the most significant gene is *Socs2*, which is a well-known signature for promoting proliferation and dampening the differentiation process^49–51^. This partially explains the huge gap between proliferation and differentiation rates for meta-clone 5 and the resulting population sizes. Other stemness-related marker genes including *Ly6a* and *Dlk1* are identified as well^52,53^ (Fig. 5f).

Cell types are often defined based on their potential to differentiate into other cell types, for example via in vitro assays, as it was historically the case for intermediate haematopoietic progenitors^54,55^. Nevertheless, we saw with our analysis that meta-clones with similar fate realisations can produce progeny in different proportions. For example, meta-clones 2 & 7 of the human cord blood data are both multi-potent, but have different offspring sizes (Supplementary Table S5). Similarly, meta-clones 0 ∼4 of the mouse hematopoietic dataset can generate similar terminal states, but each produces mainly a certain population, as shown in Fig. 5b and Supplementary Table S8. This analysis suggests that we can further idetify subpopulations of HSPCs based on the magnitude of their produced progeny.

Similarly, the DEGs of meta-clones 1∼4 compared to meta-clone 0 (Supplementary Fig. S17 and S18) illustrate that, for multi-potent clones, the transcriptomic differences in early stages can predict the population size of the offspring. Specifically, the volcano plot and stacked violin plot for the comparison of meta-clones 4 and 0 reveals a few genes of interest (Fig. 5g, 5h). Compared with meta-clone 0 (which is multi-potent but more biased towards the Monocyte lineage), meta-clone 4 is also a multi-potent clone but rather biased towards erythrocytes, mast cells and megakaryocytes. Significant genes include *Itga2b*, which is specifically up-regulated during megakaryopoiesis^56,57^, and *Gzmb*, which is highly expressed in mast cells and used to degrade hemidesmosomal proteins^58,59^. Interestingly, *Akr1c13* and *Hbb-bt* were also found to be significantly expressed (Fig. 5i). *Akr1c13* is highly expressed in the Basophil population although this is not a dominant population in meta-clone 4 (meta-clone 2 is dominant in the Basophil population). However, this in consistent with the differential cell counts: indeed, meta-clone 4 produces a larger Basophil population compared with meta-clone 0 (Fig. 5i, left, Supplementary Table S8). Moreover, gene *Hbb-bt* is a mutated variant of the *β*-globin genes, and can cause an ineffective erythropoiesis process and lead to the apoptosis process of erythroid precursors^60–62^. This might explain why there are 11,885 prog_Meg_Ery cells in total in meta-clone 4, whilst the number of produced Megakaryocytes and Erythroids is roughly 3:1. We thus advocate for a redefined definition of cell types or sub-types that includes, besides the fate-bias, the size of the generated progenies.

In conclusion, based on the transcriptomic signature clone, we could predict the lineage preferences as well as the population size of terminal states of meta-clone 4. This further strengthens the hypothesis that cell fate is initiated at very early stages.

## Discussion

In this paper, we presented CLADES, a NeuralODE-based method to reconstruct and interpolate total cell counts along a differentiation time course. With the help of the Gillespie algorithm, CLADES delineates detailed division topology and summaries the fate bias of progenitor cells.

A bootstrapping approach was implemented so that CLADES can robustly compare the differences between meta-clones. We also demonstrated that CLADES has better prediction results in terms of fate bias on meta-clone level compared with the existing method CoSpar, with associated quantitative measurement of meta-clones’ preference of lineages and kinetic rates. This may bring us new insights into defining a cluster not only based on ‘what’ it is producing but also on ‘how much’ it is producing.

Many works in the field of mathematical modeling for hematopoiesis have assumed that adult physiological hematopoiesis is in perfectly homeostatic conditions (referred to as “steady state”), where transition rates are constant and so are the populations’ sizes. On the other hand, more recent models^33,63^ have started incorporating the idea that, given that the number of HCSs increases over time and the relative abundance of the different populations varies upon aging, there is no steady state. CLADES can partially address this issue thanks to the constant vs dynamic mode option. Indeed, if the constant mode outperforms the dynamic one in a given system, we can conclude that the kinetic rates are nearly constant, which may lead to a stationary growth or a steady state, while if the dynamical mode performs better, the rates are time dependent and it is more likely that the system is not, and will not reach, a steady state.

Though CLADES has offered new perspectives to explore the LT-scSeq data, there are still a few challenges and limitations which need to be resolved in future works. Firstly, CLADES aims to quantitatively summarise the differentiation kinetics and fate bias of each clone as the primary goal, while examining the regulation and determination from the cellular transcriptomes in a separate step. Future work includes combining both barcodes and transcription data to infer transition rates and fate decision bias of clones, and mapping cells without barcoding information to meta-clones, since limited barcoding efficiency can affect the performance of algorithm and DEG analysis as well. Secondly, CLADES is formulated based on static barcoding with viral integration and multiple time points design. Therefore it cannot be used to analyse data from cumulative barcoding techniques, like CRISPR-Cas9 DNA editing^20,64^, which allows multiple-time barcoding and hence provides a fine-grained structure of sub-clones. However, for retrospective barcoding (often with endogenous genetic variants, for instance, MAESTER^65^), we expect that CLADES can be applied with minimal extension. Therefore it could serve as a promising tool for analyzing LT-scSeq data and continuing to make contributions to the general lineage tracing field.

Finally, although we focused on the hematopoietic system in both human and mouse to evaluate our model and revealed a few biological insights, CLADES is in principle broadly applicable to analyze other development systems. Moreover, it may be further applied to study cancer progression where different clones may have distinct phenotypic properties, e.g., cancer plasticity^66^.

## Methods

### Definitions and data structures

Denote the number of time points available as *T*, the number of meta-clones as *C* and the number of populations as *P* . We define the following terms for each individual meta-clone *c*,

- *x*_*t*,*p*_ ∈ ℝ^*T ∗P*^ : original real number of cells in the dish for population *p* and time *t*;
- *y*_*t*,*p*_ ∈ ℝ^*T ∗P*^ : number of cells sequenced from the dish for population *p* and time *t*;
- *K*_1_ ∈ ℝ^*P ∗P*^ : transition matrix of cell differentiation rates between populations. Its values are constrained to be non-negative and strictly upper triangular (to avoid the reversed differentiation process);
- *K*_2_ ∈ ℝ^1*∗P*^ : rates of the overall effects of proliferation and apoptosis process combined within a population, diagonal of the topology graph *L*;
- *L* ∈ *{*0, 1*}*^*P ∗P*^ : topology graph of cell states, binary version of *K*_1_ + *K*_2_, derived from PAGA with expert curation;
- *P*_*apop*_ ∈ *{*0, 1*}*^1*∗P*^ : vector of fully differentiated populations (terminal fates) with limited proliferation ability;
- *P*_*prol*_ ∈ *{*0, 1*}*^1*∗P*^ : vector of progenitor populations (e.g., HSCs) with strong proliferation ability;
- *µ*_*t*_ ∈ ℝ^+^: scaling factor between *x*_*t*,*p*_, *y*_*t*,*p*_ for time *t*;
- *t*_*cut*_: stabilization term used in the modified Gillespie algorithm, a trade-off between simulation accuracy and time complexity, default is 1*e*^−4^;
- *l*_*i*_: penalty terms used to regularize parameters *K*_1,2_, where *i* ∈ ℤ;
- *λ* ∈ (0, 1]: adjustable parameter controlling the magnitude of each penalty term in the loss function.

### Annotation pipeline of the human cord blood dataset

To annotate the cord blood dataset, we first transferred the label from the fetal liver atlas published by Popescu et al.^67^ by means of the Seurat label transfer algorithm (functions `FindTransferAnchors` and `TransferData`). We then clustered our landscape by means of the Leiden algorithm in scanpy package, and assigned to each cluster the cell type of the most common transferred label. Finally, we renamed the two clusters with the most immature progenitors as HSC/MPP 1 and HSC/MPP 2 in order to make sure that our differentiating hierarchy starts from the most immature cells, without contamination from differentiated progeny.

### NeuralODE based architecture

Given a population balance model, the per capita growth/transition rates (Eq. 2) can be treated as either time-invariant (constant value) or time-variant (they assume a different values at each time point). For a time-invariant scenario, the *K*_1,2_ themselves are the trainable parameters, whilst for the time-variant scenario, the ODE block is built upon a 2-layer multi-layer perceptron (MLP, the number of hidden dimensions is dependent on the number of populations, default is 32) with *x*_*t*,*p*_ as input and *K*_1,2_ as output, respectively. Softplus activation function was used since it has a unique gradient, which is theoretically better than other non-smooth nonlinear activation functions such as ReLU and LeakyReLU given the inner characteristics of NeuralODE^36^. *K*_1_ is further masked by the topology graph *L* to confine the empirically infeasible direction of transitions (e.g., backward transitions or transitions from Late Erythroids to Monocytes). Squaring ensures the inferred rates to be non-negative in *K*_1_ and the overall transition matrix *π*(*t*) can be inferred when combining the estimation rates; the `diag` function is used to transform a vector into a zero-like matrix where the diagonal is that vector,

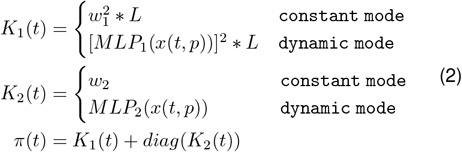

where parameter *w*_1_, *w*_2_ are weight matrix and weight vector, respectively. MLP with 2 dense layers and a relatively small hidden dimension was used because the model is run at the meta-clone rather than at the individual clone scale, which significantly reduces the number of rates to be inferred, and more layers (or dimension size) would inevitably lead to the risk of overfitting.

### Scaling factor for experimental cell counts

As scRNAseq is destructive by nature, cells sequenced at later time points cannot quantitatively reflect the accurate dynamics of cell counts during this period. Since our model is based on the real number of cells at each time point, sequenced cell counts were scaled back to total cell counts in the culturing environment based on additional information (e.g., either manual counts or data from fluorescence-activated cell sorting, FACS^68^) before being fed to the model.

In order to calculate the scaling factor between sequenced counts and real total counts of the Cord Blood data used in this manuscript, the number of cells in the dish were reported at each sequencing time which provided us with the reference cell counts (Supplementary Table S3, Supplementary Table S5). Then the estimated total number of cells at each time point is acquired in a cascading way,

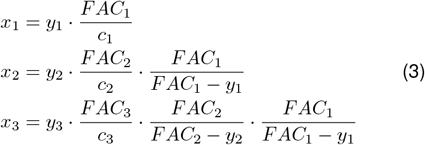

where *FAC*_1,2,3_, *y*_1,2_, *c*_1,2,3_ are the numbers of cells sorted in the dish, sequenced in the experiment and with clonal information at different time points respectively, and *x*_1,2,3_ is a cell count tensor with a shape of (time, meta-clone, population).

### Initial condition of the NeuralODE system

Solving an ODE system is essentially an initial value problem, where the first time point is usually chosen as the initial value. However, to facilitate the analysis with CLADES, we manually added an extra time point, Day 0, to the dataset, assuming that the number of initially labelled HSPCs equals the number of unique barcodes; this reduces model flexibility and estimation error.

### Parameter inference

Given a LARRY-based LT-scSeq dataset with noise due to detection or loss of barcodes, we formulate the cell counts at each time point as sampled from a Poisson distribution with means given by the neural network (we found it to be more robust than the commonly used GaussianNLL loss). We minimize the negative loglikelihood loss for each meta-clone separately as follows,

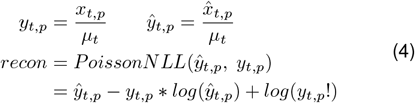

Whilst the inputs and outputs of the NeuralODE algorithm are based on real cell counts *x*_*t*,*p*_ to interpolate and mimic the natural development process, our reconstruction loss is based on the original sequenced cell counts *y*_*t*,*p*_ to avoid a stiff ODE that is difficult to solve and to speedup the back-propagation process^69^, as the number of cell counts can easily scale to millions. Besides reconstruction loss, the model also incorporates the following penalty terms as shown in Eq.(5), where the default values are *λ*_0_, *λ*_5_ = 1.0, *λ*_1_, *λ*_2_, *λ*_3_ = 0.5, and *λ*_4_ = 0.1,

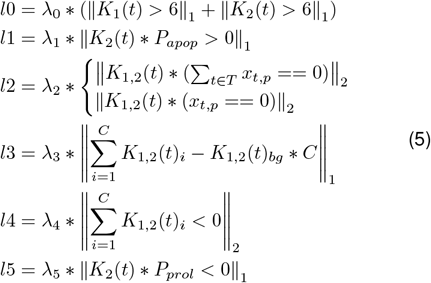

Specifically, the rational behind these penalties is:

- the transition rates in topology *L* should not be too large to violate the biological prior knowledge. Here we used 6 per day as upper bound for each rate, leaving it as an adjustable parameter at users’ discretion.
- fully differentiated populations should have limited proliferation ability (e.g., Late Erythroids).
- populations with 0 cell counts at a certain time point should not have a large proliferation or differentiation ability at that time point (cell counts are summed up in constant rate mode).
- the mean of the estimated rates for each meta-clone combined should mimic the dynamics of background cells. Here, “background” stands for all available cells, independent of the barcode presence/quality.
- we hypothesize that the apoptosis process is not too quick in a homeostatic environment.
- early progenitor populations should have an overall positive net growth rate (e.g., HSCs).

We used *L*1 norm for most penalties due to its ability of selecting non-zero parameters, except for *l*2 and *l*4, where *L*2 norm was applied to make the penalty less stringent. For *l*2, considering an ideal scenario where cell counts sequenced at each population and time point are 100% accurate, then population with 0 counts should not have either proliferation or differentiation rates. However, in reality, loss of barcodes or sequencing error at each time point could introduce extra dropouts to cell counts, especially for small-sized clones, making the data harder to analyze (e.g., in the cord blood dataset, meta-clone 1 does not have any HSC/MPP 1 or HSC/MPP 2 whilst most of the progenies exist). By using the aforementioned technique, CLADES has the ability to counteract this negative effect and automatically interpolate cell counts to recover a smoothed trajectory. For *l*4, our intention was to introduce moderate constraints to the model in terms of suppressing the negative transition rates, which would increase the model’s flexibility when facing complicated systems.

The overall cost is the sum of both penalty terms and reconstruction loss at each time point *t*, clone *c*, population *p* respectively,

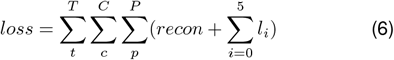

### From DNA barcodes to meta-clones

According to the experimental protocol, the inheritable DNA barcodes are induced in undifferentiated populations and their immediate progenies at Day 0. Due to the typical challenges in clonal analysis, e.g., cell dropout, barcode homoplasy or loss of barcodes^32^, although hundreds of unique clones are captured, a large proportion only contains few cells or appeared in limited time points and populations. Therefore, analyzing the behavior of individual clones is infeasible.

It is straightforward to use the pooled information, under the assumption that clones with similar kinetics should in turn produce similar cell counts at a specific time point *t* for a specific population *p*. Using cell counts at (*t*_*i*_, *p*_*i*_) as features, clones with alike characteristics were then clustered together to form meta-clones using the Leiden clustering method from scanpy package^70^. The number of meta-clones produced for each dataset is left as an adjustable parameter; in our analysis, the default way is to set the parameter `resolution` in Leiden clustering equals to 1.

### Model initialization and training details

For the constant mode, the default Kaiming Uniform^71^ was used to initialize the rates (*K*_1,2_) of the ODE function,

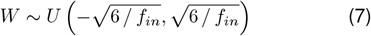

where *f*_*in*_ is the dimension of input layer (number of defined cell populations). For dynamic mode, since the estimated rates for each meta-clone are given by linear layers with a small input, the weights are relatively large if the parameters are initialized by the aforementioned approach because of the inversely proportional property. Therefore, to avoid problems like gradient exploding or numerical overflow which makes the adaptive solver unable to solve, we used standard normal distribution *N* ∼ (0, 0.01) to initialize weights in MLP alternatively.

To train the model, the default epochs are 1,500 whilst for the ODE block of dynamic mode, softplus activation and hidden dimension size of 32 were used. AdamW optimizer with default settings were adopted and we varied the learning rate using multi-step learning rate decay every 200 epochs with decay rate *γ* = 0.5. The learning rate for constant mode and dynamic mode are 5*e*^−2^ and 1*e*^−3^, respectively.

### Bootstrapping for model confidence intervals

As described in the parameter inference section, PoissonNLL was used instead of GaussianNLL because it is more robust to sparse data with lineage relationships. However, we could not directly acquire the estimation error or 95% CI based on this loss function.

To generate such data and get a comprehensive analysis of the rates, we adopted the bootstrapping strategy, which randomly samples the observation data with replacement *M* times (the initial time point *t*_0_ is not sampled). Based on the central limit theorem, we could get the percentile CI for each transition rate and the quantile estimation error for the total cell counts after ranking the fitted parameters 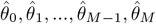 of all bootstrapping trails.

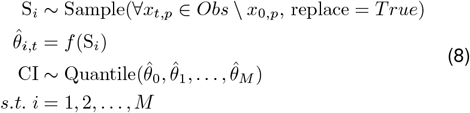

For statistical analysis and the comparison of rates within different meta-clones, we mainly used student t-test and Mann Whitney U rank test based on whether the distribution of bootstrapped values follows a standard normal distribution or not. The *p*-values of multiple tests were corrected using Benjamini-Hochberg procedure (false discovery rate, FDR-BH) with 0.05 as threshold.

The bootstrapping approach might over-estimate the CI given limited available experimental time points; alternative approaches include increasing sequencing times or profiling the likelihoods^72,73^ after fitting the model (Supplementary Note 3), at the cost of a complicated experimental design or a linearly increased computational complexity.

### Helping functions

CLADES’s performance mainly depends on the loss function and the penalties during the optimization process; additionally, we use the average recovery rate of cell counts and mean absolute difference among the rates from bootstrapping to help assessing the usage of the model.

`Average recovery rate`. This model evaluation metric is used to assess how well the model could reconstruct the original cell counts with the following mathematical formula,

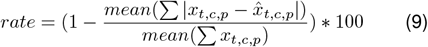

where *x*_*t*,*c*,*p*_ and 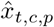 are the observed and predicted cell counts at any given time point, meta-clone id and populations.

`Mean absolute difference of parameters`. This part is mainly used to post-process the bootstrapping results together with the *p*-values given by the statistical tests. In some cases, even if the variance of distribution given by bootstrapping is small, the statistical difference among rates is still significant, which leads to the wrong interpretation especially when the total cell counts is equal to 0. Suppose we need to compare two rates and from bootstrapping we get 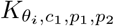 and 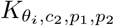 Then the mean absolute difference of rates is defined as,

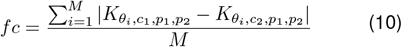

and the default threshold for this metric is 0.1: any difference lower than this threshold would be considered as not significant.

### Using Gillespie to simulate differentiation landscapes

We adopted and modified the original Gillespie algorithm to make it achieve a balance between accuracy and time complexity (Supplementary Note 2, algorithm 1). As *t*_*cut*_ was introduced in the algorithm design to guarantee convergence (e.g., the time increment between two simulation steps Δ*t* = *max*(Δ*t, t*_*cut*_)), the cell counts generated by the stochastic simulation algorithm does not perfectly resemble the observed counts, whilst this does not affect the division statistics of a progenitor cell to produce a certain progeny and fate bias towards a terminal states.

The division summary is recapitulated starting from an early progenitor cell at the initial time point (e.g., HSC/MPP 1 or prog_1); the number of proliferation events was counted until the first progeny was produced. This progeny could either be a later progenitor or a specific cell fate (e.g., prog_2 or Erythroids). In some realisations, there were only differentiation events, thus the number of division was 0.

Predicting the fate bias given a progenitor cell is a nontrivial task. Since our algorithm always starts with 1 initial progenitor cell, we run the simulation for 1,000 times to reproduce the stochasticity. Given a celltype, the fate bias prediction part of CLADES works like chained conditional probability,

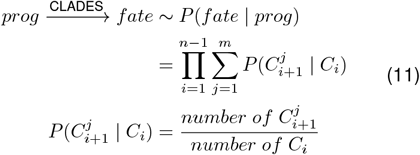

where *C*_*i*_ can be either a progenitor, terminal fate or any other intermediate cell types along the progression path and *j* denotes the different differentiation path towards the same fate (e.g., Monocytes can be produced from both prog_DC_Mono and prog_Mono).

We construct the groundtruth fate bias using clone-level information instead of cell counts. Specifically, for each meta-clone, the groundtruth from a progenitor to a certain fate is given by

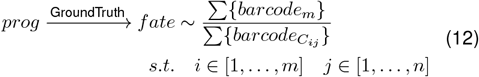

where *i* is the index of all cell types produced in a particular transition path *j* and *m* represents the fate type. Using all barcodes along the transition path within a meta-clone is an alternative strategy due to the low capturing efficiency of current LT-scSeq data, where most of clone barcodes can only be observed at limited time points or populations.

The original Gillespie algorithm sometimes falls into an infinite loop of choosing the same reaction, due to the extremely small time increments. Though the introduction of *t*_*cut*_ does not affect our mentioned analysis above, we lose other information the Gillespie algorithm could provide, e.g., analysis of cell count dynamics and statistics of reactions.

### Benchmarking with CoSpar on hematopoiesis dataset

We then applied CLADES to the same hematopoiesis dataset used by CoSpar and compared the fate transition map for all progenitor-fate pairs. We followed the standard CoSpar preprocessing pipeline described in its documentation, which provided us with bias score towards each possible cell fate named as ‘fate_map_transition_map’. Since CoSpar generates bias score for each cell and our method is based on meta-clones, to make a fair comparison, for each [meta_clone, progenitor-fate pair], we pooled the cell-specific scores of CoSpar to get a smoothed value for that meta-clone,

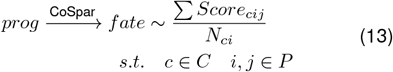

where *Score*_*cij*_ is the bias score provided by CoSpar for meta-clone *c*, from progenitor *i* to fate *j* and *N*_*ci*_ is the corresponding number of cells.

Although CoSpar provided choice of using clonal information alone to infer fate transition map, we compared with its standard form of using both lineage and gene expression information, which consistently showed better and robust performance. Fate bias scores generated by both algorithms cannot be directly compared, therefore, the relative magnitude of bias was compared after normalization, and Pearson correlation was used as a quantitative measurement,

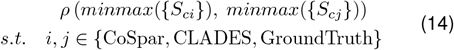

where *{S*_*ci*_*}* is the set of scores for all meta-clones given a progenitor-fate pair.

## Data availability

Two datasets were used in this paper to demonstrate the ability of CLADES: 1) the human cord blood hematopoiesis dataset is a newly generated LARRY LT-scSeq data from our collaborators which can be acquired via requests; 2) the mouse hematopoietic system is publicly available^18,32^.

## Code availability

CLADES is written in Python and is publicly available at https://github.com/StatBiomed/clonaltrans with both constant and dynamic modes implemented. It also contains documentations, pipelines and Jupyter Notebooks to reproduce figures and results mentioned in this paper.

## Acknowledgements

We kindly thank cord blood donors for their tissue donation, Dr Joanna Baxter and the team of the Cambridge Blood and Stem Cell Biobank for consenting and collecting cord blood samples. We also thank Reiner Schulte and Gabriela Grondys-Kotarba at the Cambridge Institute for Medical Research Flow Cytometry Core Facility for flow cytometry cell sorting, and the CRUK Cambridge Institute genomics centre for sequencing.

This project is supported by the National Natural Science Foundation of China (No. 62222217), Innovation Technology Commission Funding (Health@InnoHK), and the University of Hong Kong through a startup fund and a seed fund (Y.H.). This project is also funded by Wellcome (215116/Z/18/Z; E.L. and B.G.) and previous core support grants from Wellcome and Medical Research Council (MRC) to the Wellcome-MRC Cambridge Stem Cell Institute (203151/Z/16/Z). E.L. was supported by a Wellcome – Royal Society Sir Henry Dale Fellowship 107630/Z/15/Z. M.H. is supported by a Sir Henry Wellcome Postdoctoral Fellowship (224055/Z/21/Z).

This research was funded in whole, or in part, by the Wellcome Trust. For the purpose of open access, the author has applied a Creative Commons Attribution (CC BY) licence to any Author Accepted Manuscript version arising from this submission.

## Author Contributions

B.G., M.B., E.L., and Y.H. conceived the project. M.G. developed the algorithm, implemented the package CLADES, and analyzed the data with Y.H.’s and M.B.’s support. S.C., M.H. and E.C. generated the cord blood dataset. M.B. processed the data with S.C.’s and Y.C.’s support. M.G. and Y.H. wrote the manuscript with inputs from all authors.

## Competing Interests

The authors declare no competing interests.

## Notes

### Competing Interest Statement

The authors have declared no competing interest.

