## Supplementary Tables and Figures for "Unveiling Clonal Cell Fate and Differentiation Dynamics: A Hybrid NeuralODE-Gillespie Approach"

### Supplementary Information

**Table S1: Training and testing sets of a 17-day simulation data governed by ODE function with time-invariant rates.** To explore the number of training time points needed by CLADES to generate reasonable results and to test the performance in recovering the correct rates and cell counts, five independent trials were conducted using 2-6 time points as training set respectively. All the trials make use of the same five time points as test set.

| | Index | Available time points |
|---|---|---|
| Training set | 1 | Day 0, 17 |
| | 2 | Day 0, 3, 17 |
| | 3 | Day 0, 3, 10, 17 |
| | 4 | Day 0, 3, 7, 10, 17 |
| | 5 | Day 0, 3, 7, 10, 14, 17 |
| Test set | 1 | Day 2, 5, 8, 12, 15 |

**Table S2: Training and testing sets of a 8-day simulation dataset governed by ODE function with time-variant rates.** To test the performance of both constant mode and dynamic mode of CLADES on time-variant synthesized rates, four independent trials were conducted using 2-5 time points as training sets respectively. All the trials make use of the same four 4 time points as test sets.

| | Index | Available time points |
|---|---|---|
| Training set | 1 | Day 0, 8 |
| | 2 | Day 0, 4, 8 |
| | 3 | Day 0, 2, 6, 8 |
| | 4 | Day 0, 2, 4, 6, 8 |
| Test set | 1 | Day 1, 3, 5, 7 |

**Table S3: Statistics of cells and barcodes on different sequencing time points of LARRY cord blood data.** GFP+ cells with clonal barcodes are used to construct meta-clones. The number of cells at each time point were transformed back to their actual counts in culturing environment based on the total number of cells in the dish and the fraction of cells that was recultivated.

| | Day 3 | Day 10 | Day 17 |
|---|---|---|---|
| Total number of cells | 13,284 | 500,000 | 1,764,225 |
| Sequenced | 8,856 | 20,000 | 40,000 |
| Recultivated | 4,428 | 460,000 | |
| GFP+ cells with clone info | 2,482 | 2,578 | 9,073 |

**Table S4: Performance comparison of the LARRY cord blood dataset between the two modes of CLADES.** Here we mainly used two evaluation metrics: 1) spearman correlation between observed cell counts and predicted cell counts which gives us a general trend of model performance, 2) average recovery rate at each time point in percentage view, which quantitatively assesses the performance of NeuralODEs.

| | Day 3 | | Day 10 | | Day 17 | |
|---|---|---|---|---|---|---|
| | Avg. Recovery Rate | Corr | Avg. Recovery Rate | Corr | Avg. Recovery Rate | Corr |
| constant mode | 40.56% | .626 | 22.31% | .885 | 81.78% | .868 |
| dynamic mode | 56.20% | .712 | 72.57% | .889 | 91.18% | .903 |

**Table S5: Number of total cell counts from all time points combined transformed from sequenced counts of the human cord blood data.** This transformation is based on the scaling factor given by experimental design (Methods) and each meta-clone has preference not only on different lineages but also on the quantity of populations. Shaded gray region represents terminal states of the dataset.

| Populations / Meta Clone | 0 | 1 | 2 | 3 | 4 | 5 | 6 | 7 | 8 | 9 | 10 | 11 |
|---|---|---|---|---|---|---|---|---|---|---|---|---|
| HSC/MPP 1 | 15 | 0 | 50 | 0 | 5 | 50 | 15 | 556 | 0 | 0 | 0 | 0 |
| HSC/MPP 2 | 541 | 0 | 53,133 | 0 | 556 | 536 | 25 | 17,039 | 0 | 0 | 60 | 0 |
| MEMP | 48,403 | 671 | 509 | 4,266 | 10 | 0 | 0 | 1,040 | 816 | 5 | 0 | 30 |
| Mast cell | 380,089 | 32,998 | 4,590 | 35,209 | 2,121 | 0 | 6,127 | 139,773 | 27,395 | 3,666 | 0 | 2,633 |
| Early Erythroid | 274,774 | 62,422 | 0 | 104,848 | 5 | 0 | 0 | 3,074 | 3,700 | 531 | 0 | 225 |
| Mid Erythroid | 468,916 | 175,937 | 3,607 | 43,232 | 0 | 531 | 0 | 62,587 | 5,797 | 531 | 0 | 1,593 |
| Late Erythroid | 78,982 | 16,923 | 8,163 | 9,249 | 509 | 0 | 0 | 8,648 | 0 | 0 | 0 | 2,653 |
| NMP | 32,870 | 1,050 | 60,554 | 0 | 27,028 | 18,575 | 51,869 | 795,568 | 0 | 0 | 2,825 | 0 |
| Mono precursor | 8,161 | 0 | 104,431 | 509 | 6,368 | 49,326 | 13,687 | 213,723 | 0 | 0 | 35 | 0 |
| Monocyte | 64,647 | 5,089 | 66,882 | 3,054 | 88,067 | 81,257 | 213,544 | 1,181,961 | 15 | 0 | 8,959 | 0 |
| DC precursor | 1,526 | 0 | 124,221 | 0 | 0 | 6,898 | 1,017 | 509 | 0 | 0 | 0 | 0 |
| DC | 0 | 0 | 96,954 | | 1,017 | 6,370 | 1,017 | 2,609 | 0 | 0 | 0 | 0 |

**Table S6: Estimated statistics of cells and barcodes on different sequencing time points of mouse hematopoietic data.** Cells with clonal barcodes are used to construct meta-clones. Since the original experiment did not capture the exact cell counts, we used estimated counts based on the total expansion of the culture being roughly (4-6)-fold in the first two days, (4-6)-fold between days 2 and 4, and (2-4)-fold between days 4-6. The initial number of cells is set to 5,859 because there were 5,859 unique barcodes.

| | Day 0 | Day 2 | Day 4 | Day 6 |
|---|---|---|---|---|
| Estimated total number of cells | 6,000 | 30,000 | 150,000 | 450,000 |
| Sequenced | | 27,757 | 47,444 | 51,660 |
| Cells with clone info | | 4,558 | 14,679 | 28,742 |

**Table S7: Performance comparison of the mouse hematopoietic dataset between constant and dynamic modes of CLADES.** As shown in the table, two modes have similar performance based on evaluation metrics, indicating that the dynamic behavior of this system is not so complex.

| | Day 2 | | Day 4 | | Day 6 | |
|---|---|---|---|---|---|---|
| | Avg. Recovery Rate | Corr | Avg. Recovery Rate | Corr | Avg. Recovery Rate | Corr |
| constant mode | 45.52% | .760 | 74.08% | .917 | 83.25% | .934 |
| dynamic mode | 53.33% | .775 | 78.25% | .907 | 85.68% | .935 |

**Table S8: Number of total cell counts summed from all time points of the mouse hematopoietic dataset.** The scaling factor for transformation is derived from the fold change of total expansion and each meta-clone has preference not only on different lineages but also on the magnitude of the produced populations. Shaded gray region represents terminal states of the dataset. Upper table, estimated total cell counts. Lower table, original sequenced cell counts.

| Populatoins / Meta Clone | 0 | 1 | 2 | 3 | 4 | 5 | 6 | 7 | 8 | 9 | 10 | 11 | 12 |
|---|---|---|---|---|---|---|---|---|---|---|---|---|---|
| prog_1 | 867 | 962 | 602 | 508 | 773 | 4,308 | 468 | 286 | 1,902 | 201 | 115 | 35 | 15 |
| prog_2 | 3,539 | 5,630 | 830 | 394 | 556 | 4,789 | 636 | 191 | 13,728 | 92 | 13 | 0 | 89 |
| prog_3 | 7,803 | 1,004 | 108 | 227 | 5 | 360 | 191 | 368 | 464 | 182 | 3 | 3 | 0 |
| prog_4 | 3,687 | 17,024 | 993 | 1,430 | 174 | 1,587 | 1,518 | 182 | 1,961 | 16 | 35 | 1 | 22 |
| prog_Baso_Meg_Ery_Mast | 41 | 1,515 | 5,782 | 27 | 844 | 263 | 142 | 0 | 829 | 1 | 204 | 0 | 0 |
| prog_Baso_Eos | 101 | 728 | 7,928 | 87 | 315 | 23 | 46 | 11 | 8 | 0 | 177 | 0 | 3 |
| prog_DC_Mono | 422 | 40 | 21 | 14 | 11 | 18 | 11 | 153 | 0 | 936 | 0 | 0 | 0 |
| prog_Ly_pDC | 719 | 41 | 40 | 1 | 32 | 31 | 85 | 61 | 15 | 1,250 | 0 | 0 | 0 |
| prog_Meg_Ery | 27 | 120 | 904 | 53 | 11,885 | 155 | 36 | 0 | 501 | 11 | 291 | 0 | 0 |
| prog_Mono | 32,782 | 1,758 | 200 | 1,247 | 9 | 126 | 222 | 3,129 | 60 | 99 | 9 | 128 | 0 |
| early_prog_Neu | 4,235 | 17,678 | 842 | 4,663 | 55 | 95 | 207 | 310 | 29 | 0 | 23 | 0 | 3 |
| prog_Neu | 729 | 1,604 | 812 | 11,578 | 29 | 0 | 187 | 372 | 0 | 0 | 0 | 3 | 0 |
| Baso | 63 | 188 | 28,700 | 282 | 497 | 0 | 52 | 0 | 0 | 0 | 1,224 | 0 | 6 |
| DC | 113 | 12 | 29 | 11 | 0 | 0 | 3 | 77 | 0 | 133 | 0 | 0 | 0 |
| Eos | 3 | 26 | 1,100 | 159 | 34 | 3 | 37 | 0 | 0 | 0 | 17 | 0 | 3 |
| Ery | 8 | 0 | 91 | 9 | 2,459 | 0 | 0 | 0 | 0 | 0 | 40 | 0 | 0 |
| Ly | 190 | 0 | 0 | 0 | 0 | 3 | 23 | 20 | 23 | 627 | 0 | 0 | 0 |
| Mast | 46 | 46 | 1,013 | 99 | 1,060 | 9 | 0 | 23 | 0 | 0 | 9,815 | 0 | 0 |
| Meg | 17 | 20 | 223 | 36 | 6,490 | 0 | 3 | 0 | 26 | 0 | 37 | 0 | 0 |
| Mono | 14,781 | 308 | 130 | 3,077 | 63 | 26 | 64 | 4,871 | 8 | 99 | 17 | 22 | 0 |
| Neu | 145 | 63 | 412 | 12,809 | 34 | 0 | 113 | 431 | 0 | 0 | 0 | 0 | 0 |
| pDC | 8 | 0 | 34 | 0 | 0 | 0 | 0 | 0 | 0 | 180 | 0 | 0 | 0 |

| Populatoins / Meta Clone | 0 | 1 | 2 | 3 | 4 | 5 | 6 | 7 | 8 | 9 | 10 | 11 | 12 |
|---|---|---|---|---|---|---|---|---|---|---|---|---|---|
| prog_1 | 761 | 786 | 568 | 508 | 509 | 1,063 | 432 | 286 | 543 | 188 | 115 | 35 | 15 |
| prog_2 | 835 | 1,143 | 237 | 290 | 117 | 798 | 403 | 145 | 1,694 | 23 | 5 | 0 | 69 |
| prog_3 | 1,427 | 163 | 26 | 78 | 3 | 50 | 60 | 191 | 56 | 33 | 1 | 3 | 0 |
| prog_4 | 602 | 2,367 | 169 | 560 | 26 | 193 | 432 | 67 | 231 | 5 | 6 | 1 | 5 |
| prog_Baso_Meg_Ery_Mast | 7 | 195 | 962 | 10 | 124 | 34 | 32 | 0 | 101 | 1 | 48 | 0 | 0 |
| prog_Baso_Eos | 13 | 85 | 1,211 | 17 | 41 | 4 | 8 | 2 | 1 | 0 | 32 | 0 | 1 |
| prog_DC_Mono | 61 | 6 | 12 | 3 | 2 | 3 | 2 | 31 | 0 | 137 | 0 | 0 | 0 |
| prog_Ly_pDC | 196 | 18 | 11 | 1 | 11 | 14 | 70 | 34 | 6 | 399 | 0 | 0 | 0 |
| prog_Meg_Ery | 8 | 22 | 171 | 11 | 1,992 | 20 | 11 | 0 | 64 | 2 | 71 | 0 | 0 |
| prog_Mono | 3,975 | 220 | 29 | 269 | 2 | 18 | 32 | 971 | 7 | 14 | 3 | 119 | 0 |
| early_prog_Neu | 527 | 2,090 | 118 | 1,314 | 7 | 11 | 43 | 84 | 4 | 0 | 4 | 0 | 1 |
| prog_Neu | 87 | 197 | 95 | 1,737 | 4 | 0 | 26 | 53 | 0 | 0 | 0 | 1 | 0 |
| Baso | 8 | 23 | 3,802 | 34 | 61 | 0 | 6 | 0 | 0 | 0 | 161 | 0 | 2 |
| DC | 13 | 3 | 6 | 2 | 0 | 0 | 1 | 14 | 0 | 16 | 0 | 0 | 0 |
| Eos | 1 | 3 | 148 | 19 | 4 | 1 | 5 | 0 | 0 | 0 | 2 | 0 | 1 |
| Ery | 1 | 0 | 12 | 2 | 324 | 0 | 0 | 0 | 0 | 0 | 6 | 0 | 0 |
| Ly | 27 | 0 | 0 | 0 | 0 | 1 | 4 | 3 | 4 | 116 | 0 | 0 | 0 |
| Mast | 6 | 6 | 125 | 14 | 130 | 2 | 0 | 4 | 0 | 0 | 1,300 | 0 | 0 |
| Meg | 2 | 3 | 27 | 7 | 899 | 0 | 1 | 0 | 3 | 0 | 5 | 0 | 0 |
| Mono | 1,757 | 38 | 20 | 384 | 8 | 3 | 10 | 881 | 1 | 14 | 2 | 7 | 0 |
| Neu | 18 | 8 | 48 | 1,584 | 4 | 0 | 13 | 54 | 0 | 0 | 0 | 0 | 0 |
| pDC | 1 | 0 | 4 | 0 | 0 | 0 | 0 | 0 | 0 | 38 | 0 | 0 | 0 |

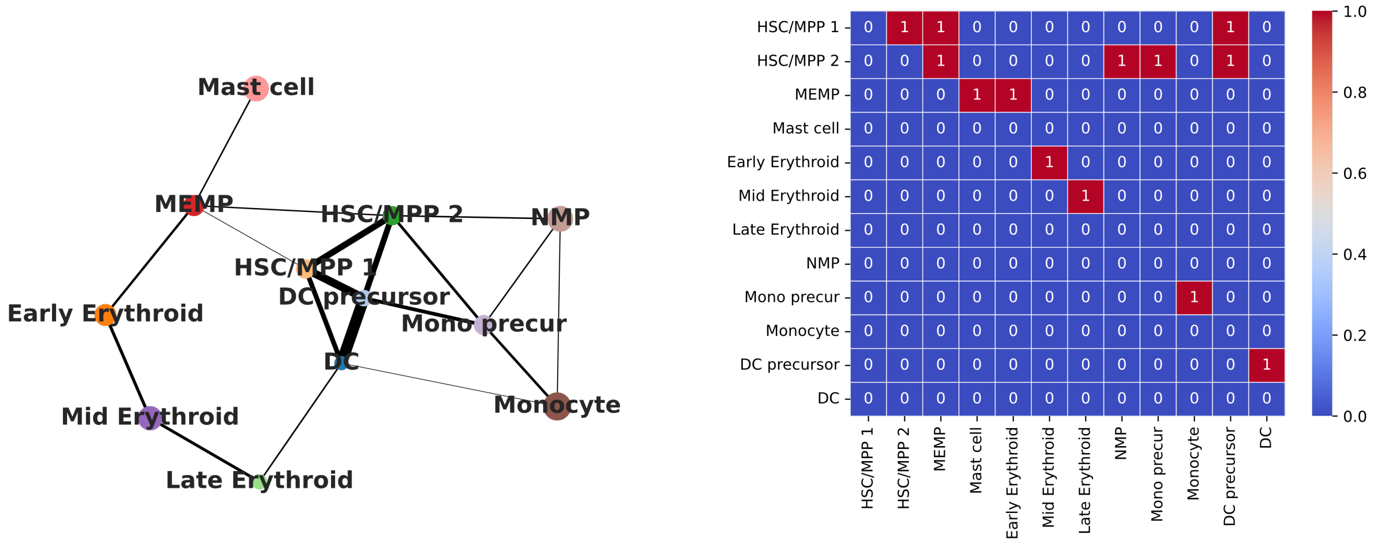


**Figure S1: Example of PAGA graph and its associated binary transition directions graph $L$.** Left, CLADES takes PAGA graph with expert curation as input, which indicates putative transition directions between populations. Right, PAGA graph is converted into a binary matrix, where 1 indicates that the population for a certain row could differentiate into the population at the corresponding column.

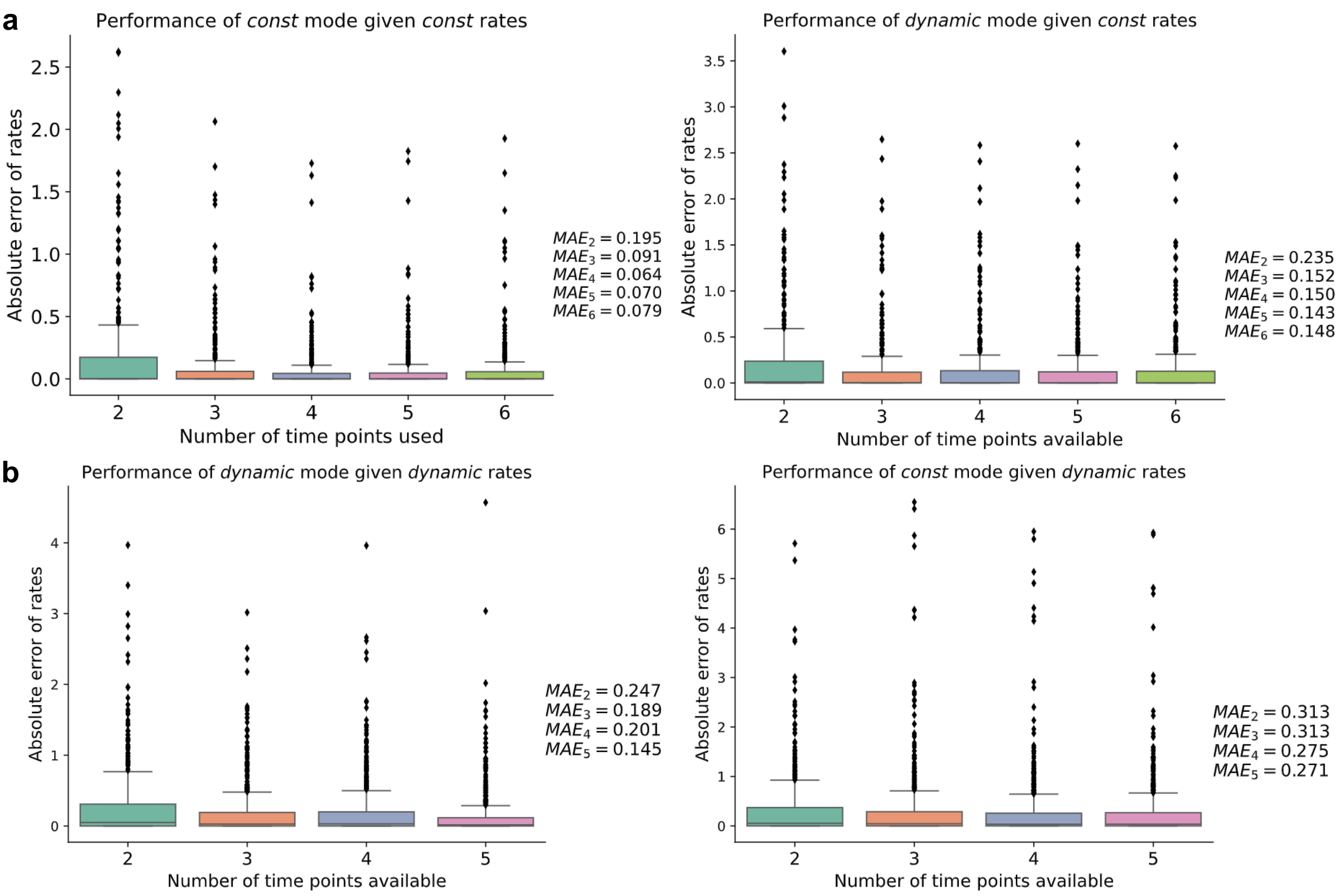


**Figure S2: Absolute error of transition rates between estimated values and the ground-truth on all simulation datasets.** We found that both constant and dynamic mode work better than the other if applied on data generated by the corresponding rates. However, the performance of the dynamic mode generally increases and is better than the constant mode when more training data are given or facing complex scenarios, suggesting the dynamic mode is more flexible to different situations.

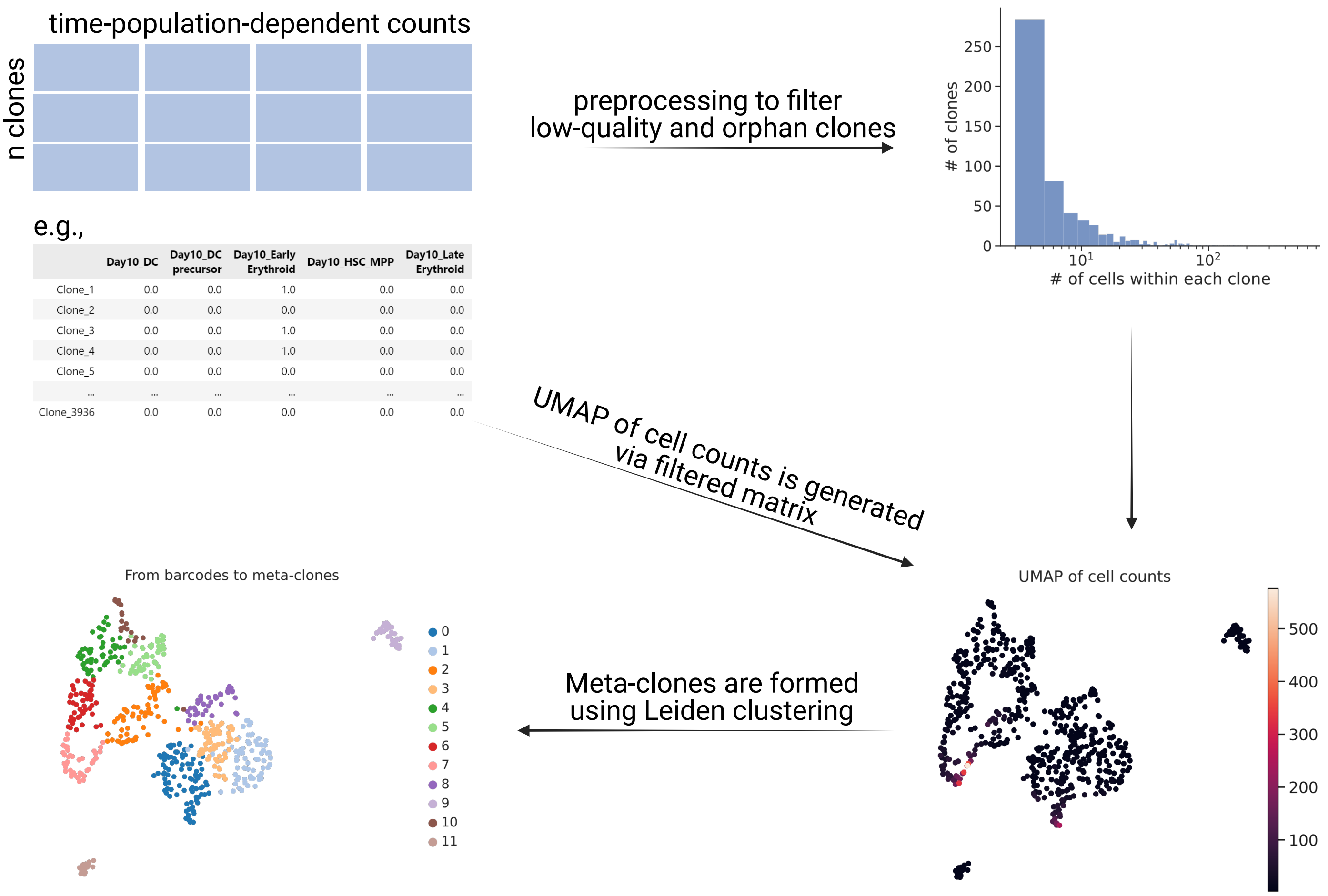


| | Day10_DC | Day10_DC precursor | Day10_Early Erythroid | Day10_HSC_MPP | Day10_Late Erythroid |
|---|---|---|---|---|---|
| Clone_1 | 0.0 | 0.0 | 1.0 | 0.0 | 0.0 |
| Clone_2 | 0.0 | 0.0 | 0.0 | 0.0 | 0.0 |
| Clone_3 | 0.0 | 0.0 | 1.0 | 0.0 | 0.0 |
| Clone_4 | 0.0 | 0.0 | 1.0 | 0.0 | 0.0 |
| Clone_5 | 0.0 | 0.0 | 0.0 | 0.0 | 0.0 |
| ... | ... | ... | ... | ... | ... |
| Clone_3936 | 0.0 | 0.0 | 0.0 | 0.0 | 0.0 |

**Figure S3: From individual DNA barcodes to the aggregated meta-clones.** Based on the barcoding information and defined populations, we could get a cell counts matrix describing characteristics of each clone. Since most of the clones are low-quality clones, containing only a few cells along the entire time-course, they were further filtered with 3 cells as a minimum threshold. The the embeddings were generated based on the time-population-dependent cell counts. Finally, meta-clones were created using leiden clustering with columns as features where each dot in the UMAP is an original clone. All the subsequent analysis and algorithms are based on meta-clone scale.

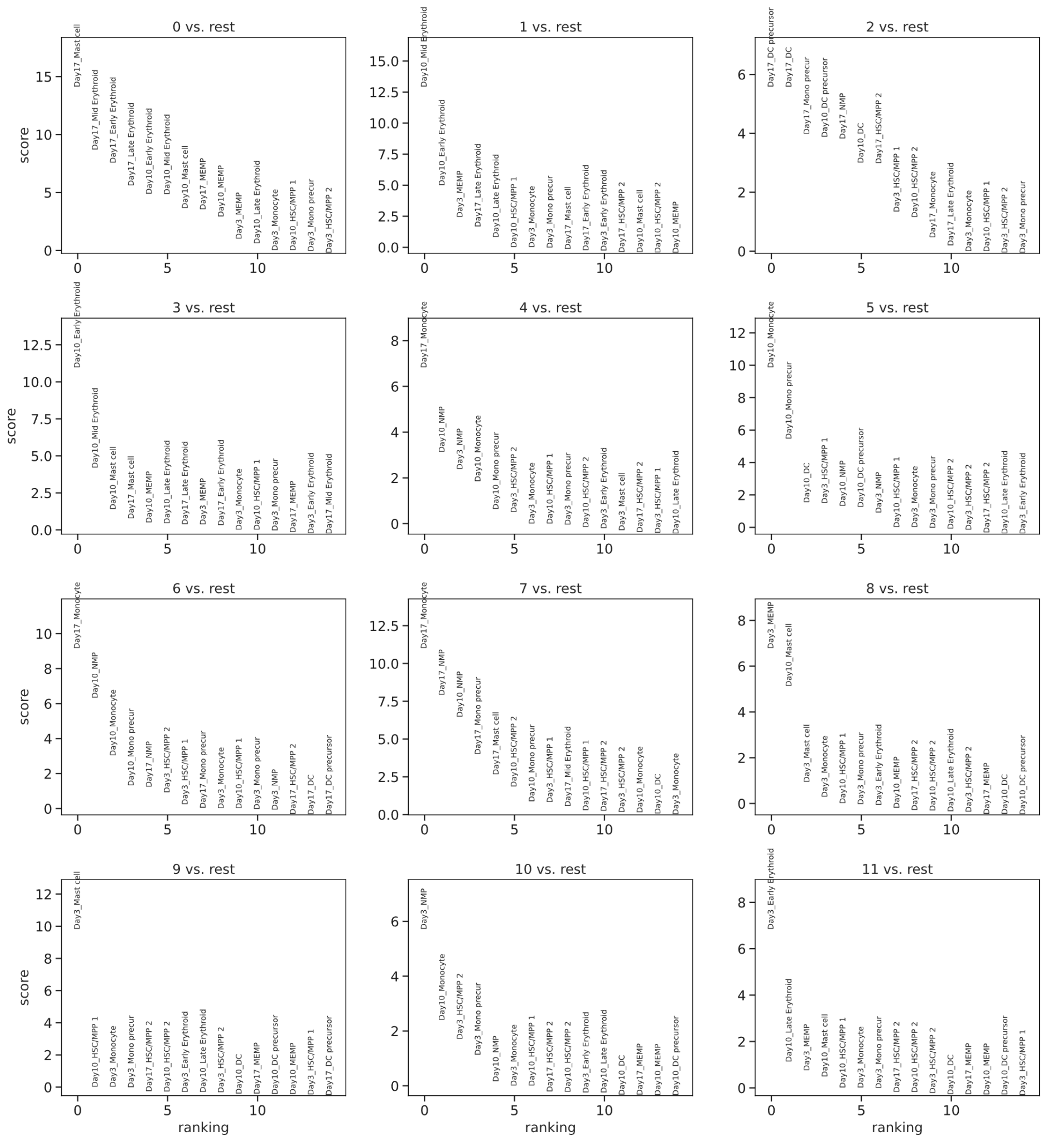


**Figure S4: From individual DNA barcodes to the aggregated meta-clones.** We used the Leiden clustering algorithm from the scanpy package with default parameters to construct meta-clones, which results in 12 meta-clones in total. Clustering features are the time-population-dependent of cell counts.

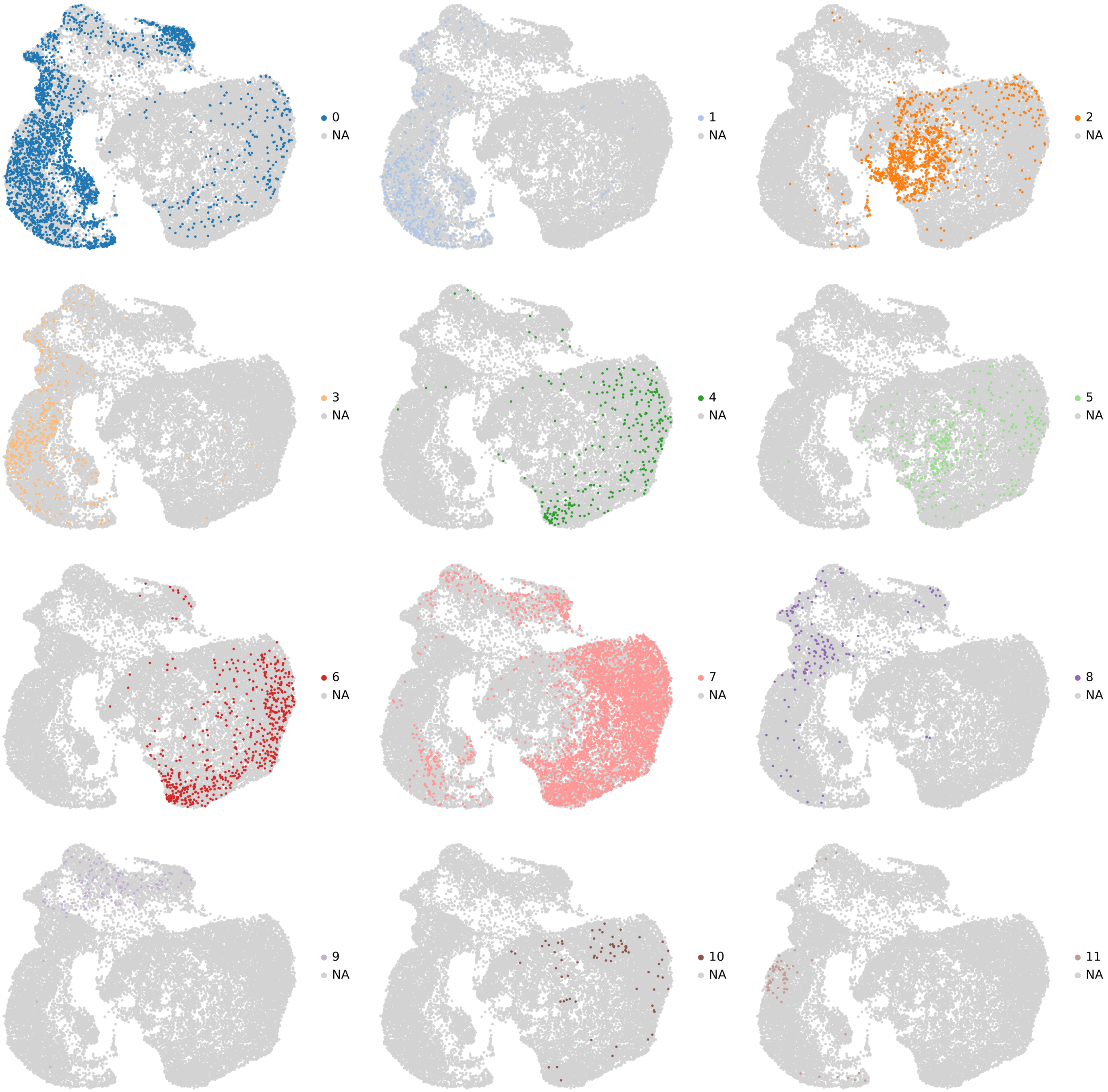


**Figure S5: UMAP illustration for each meta-clone.** Meta-clones have distinct differentiation preferences and, consequently, terminal fate states.

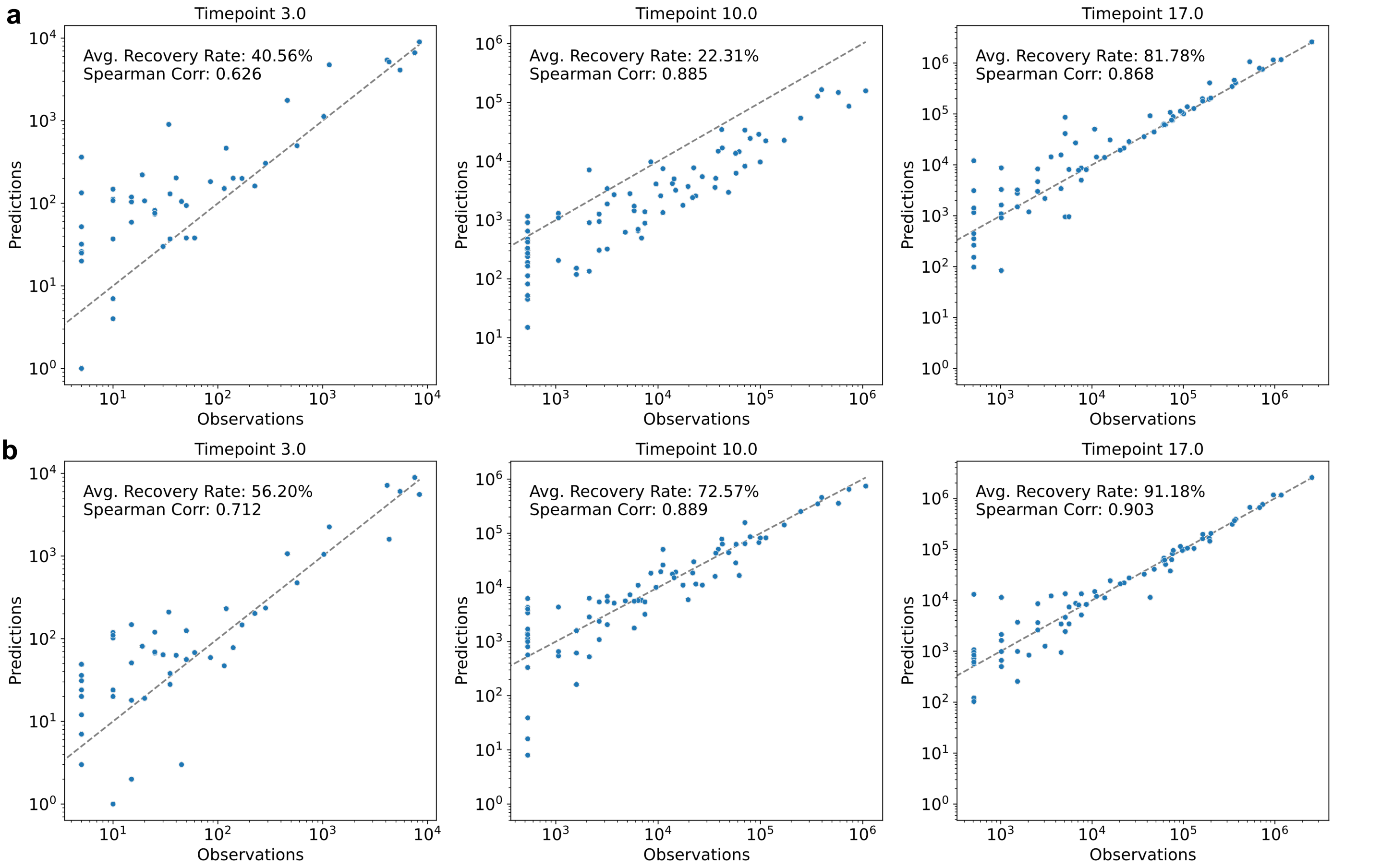


**Figure S6: Reconstruction results of CLADES on a LARRY dataset of human cordblood hematopoiesis. a,** Performance of constant mode on sequencing time points. **b,** Performance of dynamic mode on sequencing time points. The dyanmic mode exceeds constant mode in both predicting cell counts and correlation analysis, suggesting its flexibility and accuracy.

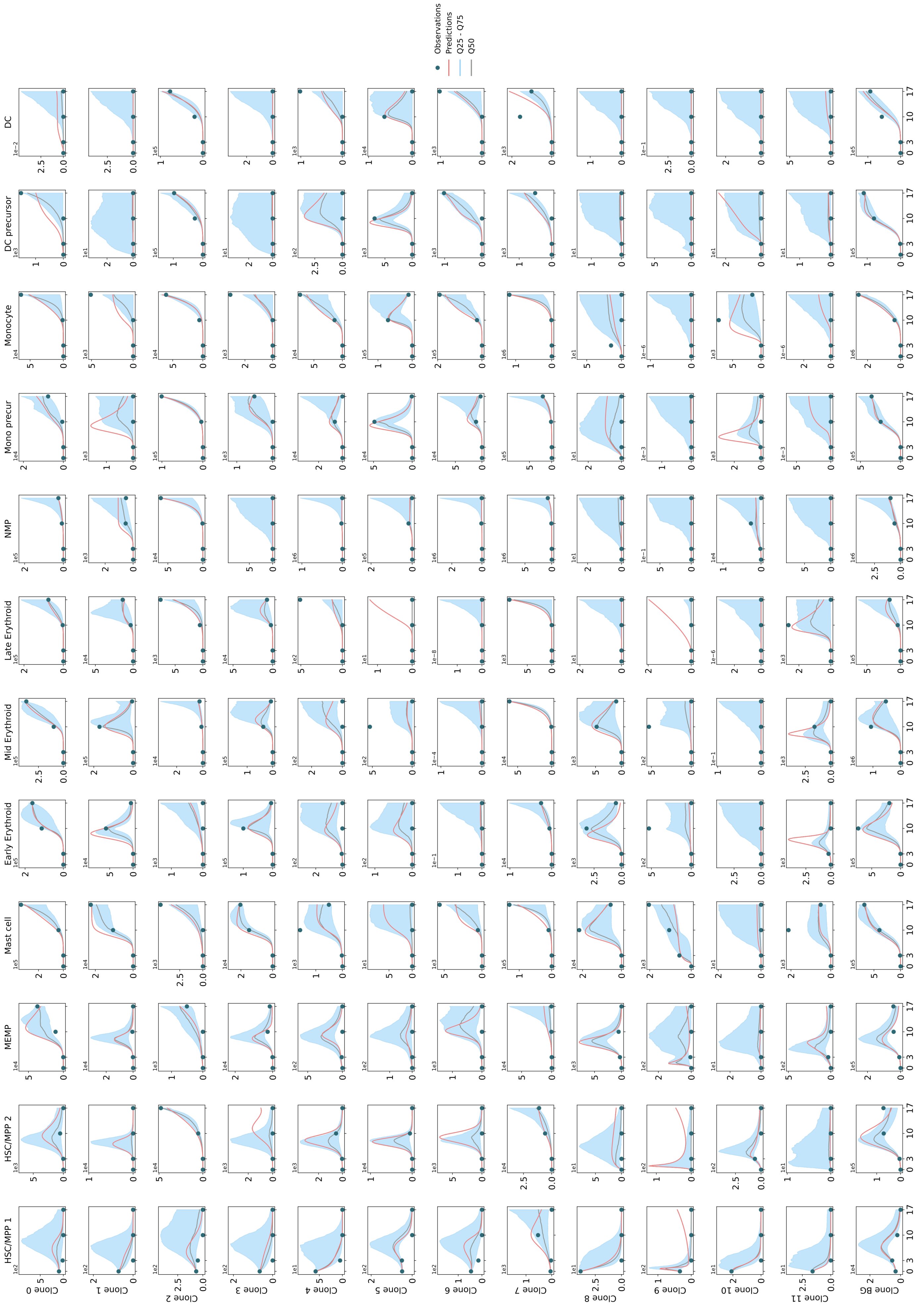


**Figure S7: Interpolation results of the dynamic mode of CLADES on human cordblood dataset along the entire time course.** The dynamic mode generally captured the trajectories well; green dots are the observed data, red line is the model fit, grey line is the median of all bootstrapping trials and the shaded light blue represents the region between the $Q25$~$Q75$ quantiles.

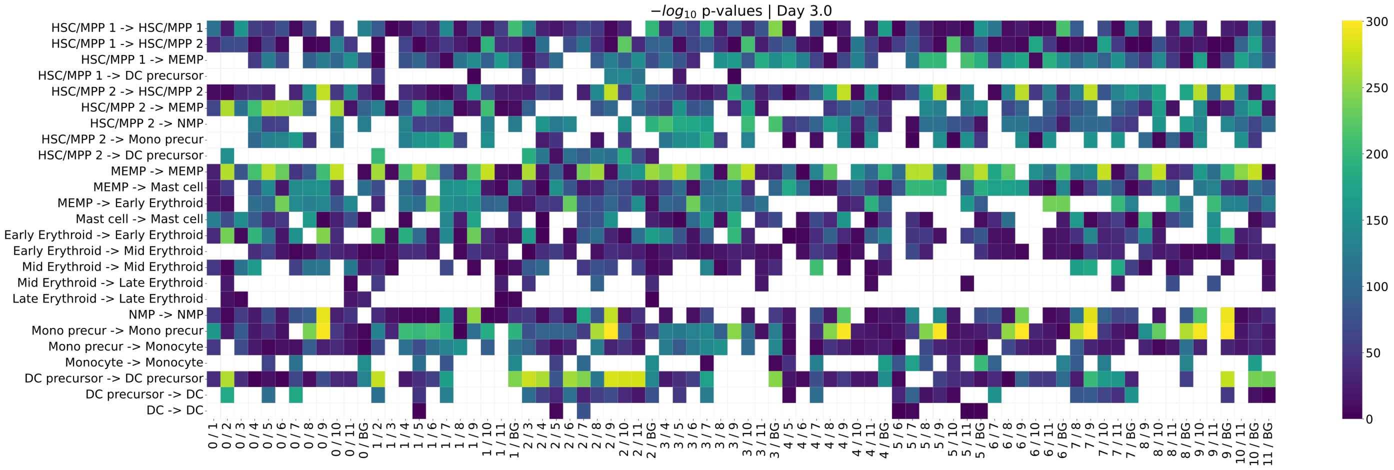


**Figure S8: Statistical tests confirms the heterogeneity of transition rates among the meta-clones and with respect to the background.** The student t-test and Mann-Whitney U rank test were used to determine whether the bootstrapping derived rate distributions are significantly different from each pairwise meta-clones. We also used the mean absolute difference of all bootstrapping trials to further filter outliers. The heatmap uses $-log_{10}(p)$ value for better visualization purpose, therefore, lighter entries represent bigger differences between a specific pair of meta-clones. Blank indicates no significance or the mean absolute difference has not passed the filtering criteria (which is an adjustable parameter).

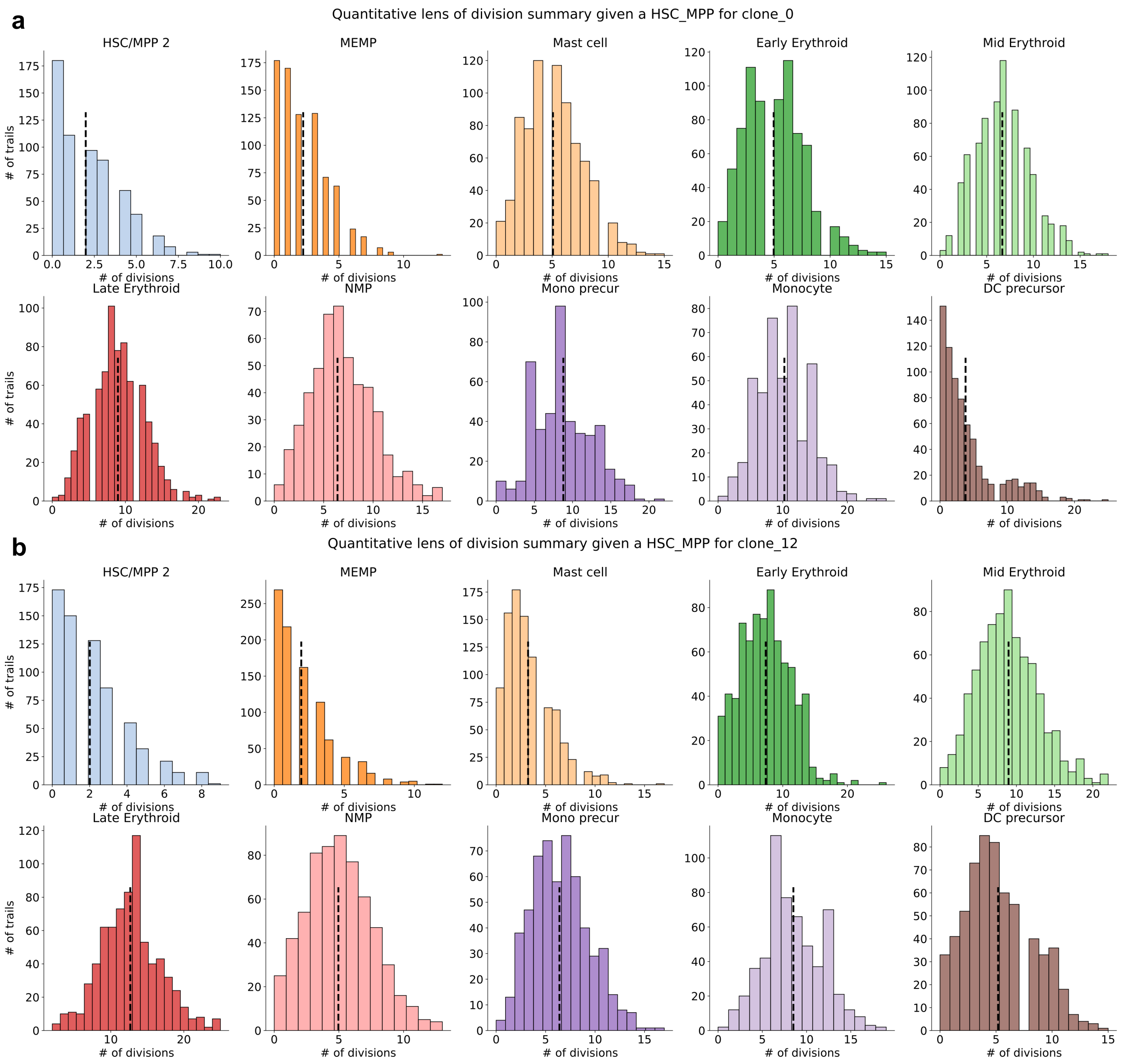


**Figure S9: Summary of division events for meta-clone 0 (a) and metaclone 12 (b, the 'background' in the main text) of human cord blood dataset given a HSC_MPP 1 as initial cell.** The histogram above shows number of division events needed to produce the first progeny given a HSC/MPP 1 as initial condition, where the dotted line indicates the mean value of the distribution. Thus, we could have a quantitative view of the characteristics of each meta-clone and the differences compared with the system.

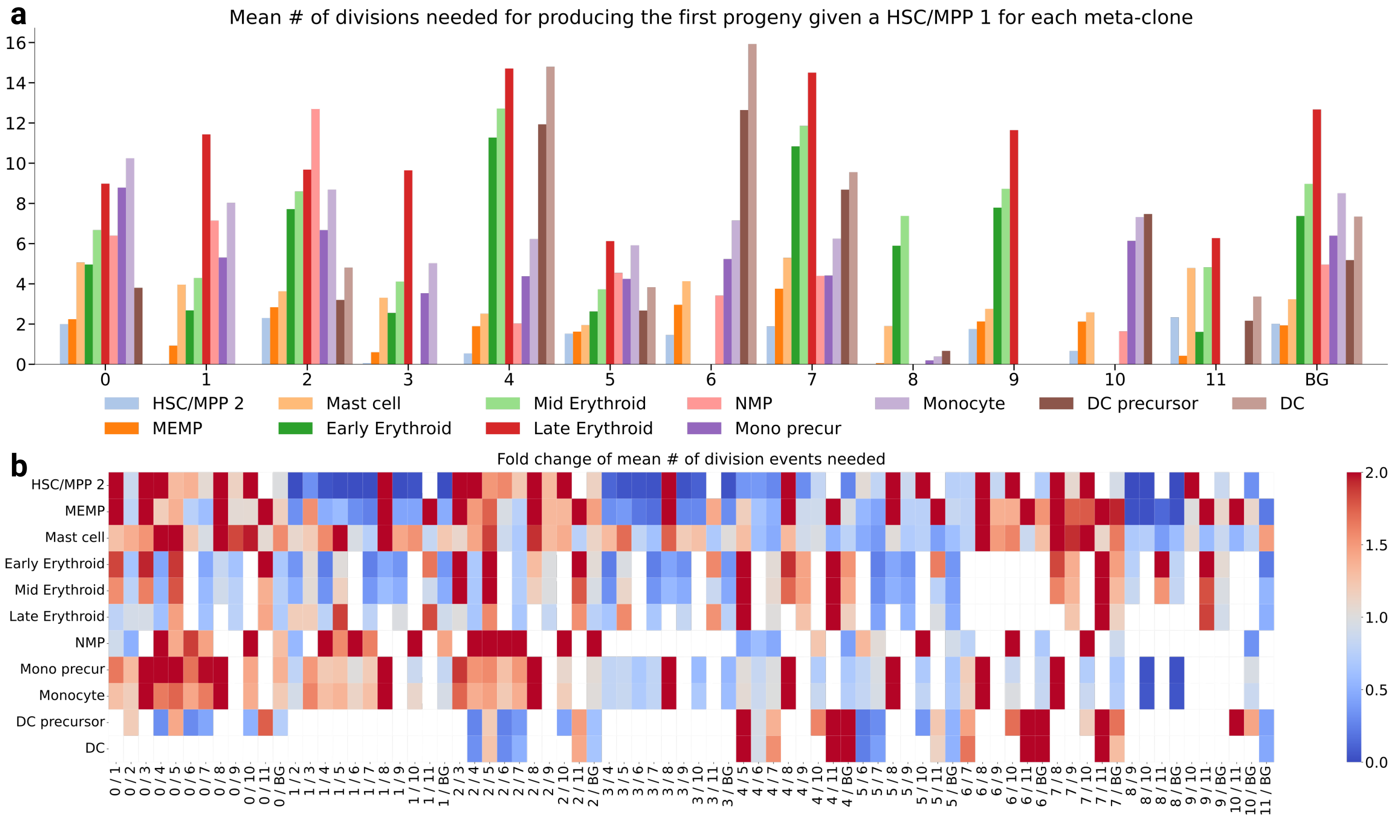


**Figure S10: Statistical tests confirms the heterogeneity of division summary among meta-clones and with respect to the background. a,** Overall result of the mean number of division events needed to produce the first progeny for each meta-clones. **b,** We used the fold change of the mean number of division events needed to produce the first progeny given a HSC/MPP 1 to illustrate the difference among meta-clones and background cells. Red indicates enhancement, blue represents smaller number of divisions and the range of color bar is (0, 2) for illustration purpose (the actual upper bound of the fold change is larger than 2). Blank region means no significance or the meta-clone does not produce the corresponding population.

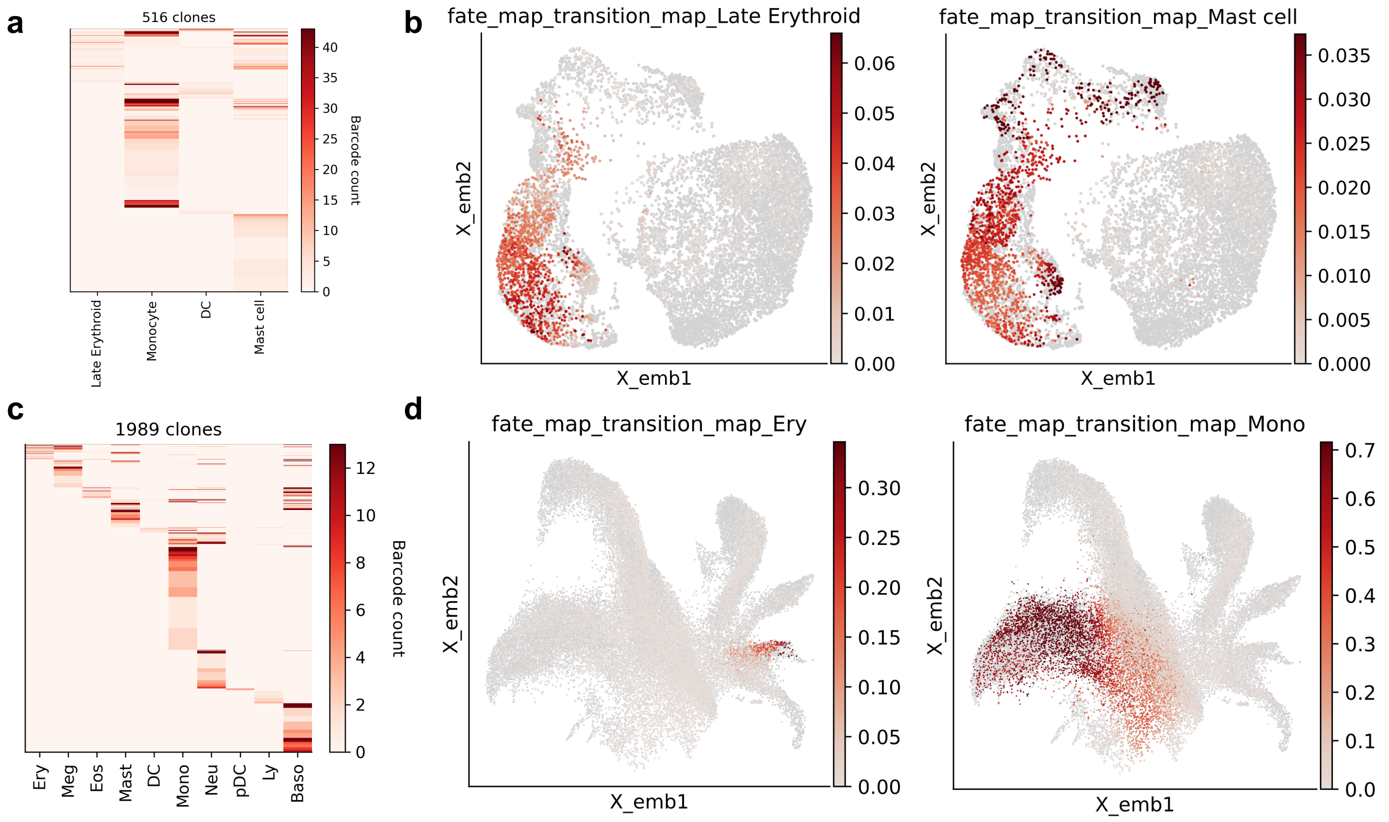


**Figure S11: Illustration of actual fate bias for each individual clone and single-cell resolution fate bias prediction of CoSpar. a,** Distribution of terminal states of each individual clone in human cord blood. Color denotes the number of cells in each clone (y-axis) for a certain terminal state (x-axis). **b,** Example of predicted fate bias score for Late Erythroid and Mast cell lineage in single-cell resolution, with higher score indicating higher bias. **c,** Distribution of terminal states of each individual clone in mouse hematopoietic system. Color denotes the number of cells in each clone (y-axis) for a certain terminal state (x-axis). **d,** Example of predicted fate biase score for Ery and Mono lineages for the mouse dataset.

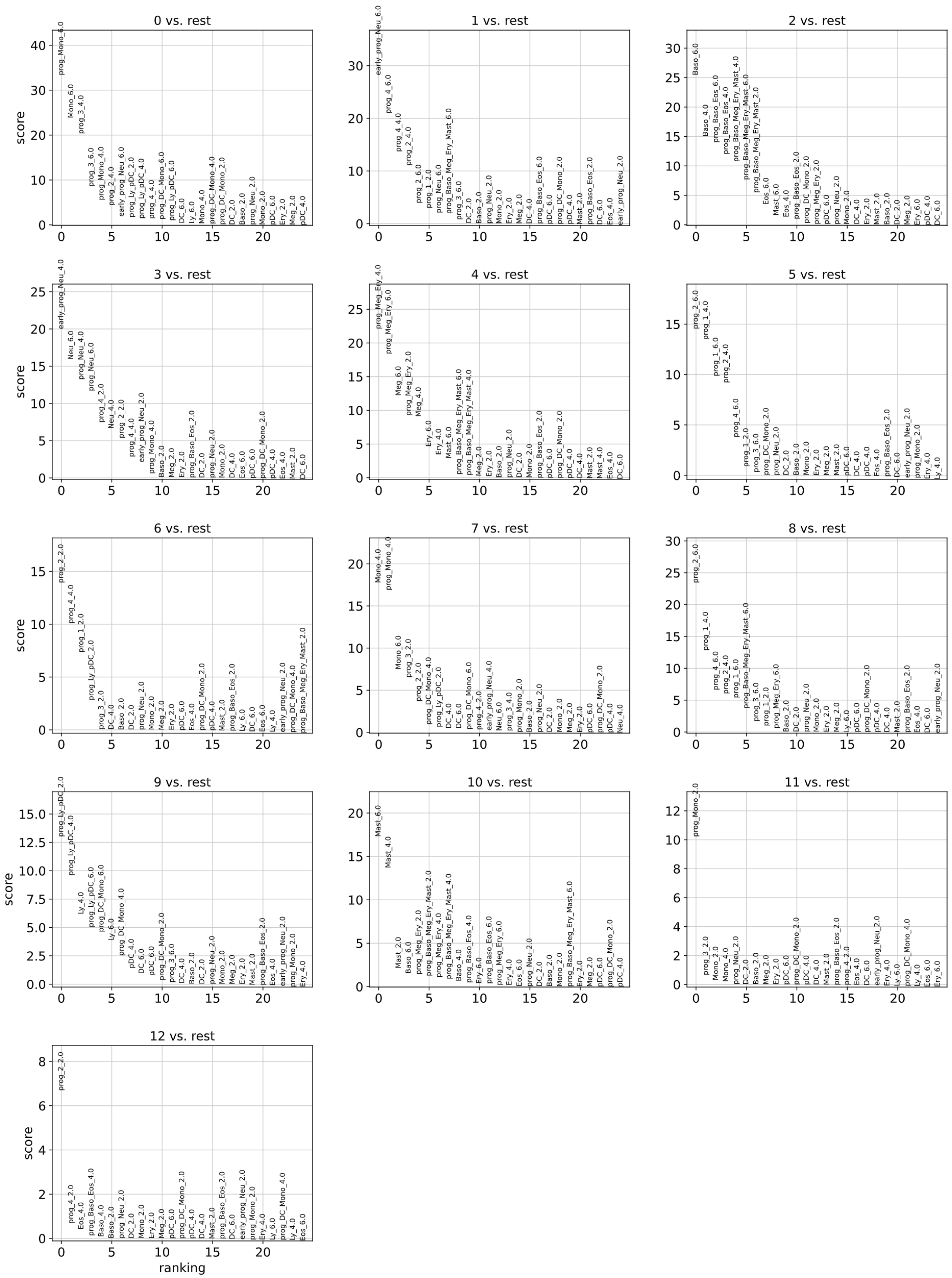


**Figure S12: Construction of meta-clones for mouse hematopoietic system.** We used the Leiden clustering algorithm from scanpy package to construct meta-clones with default parameter, which results in 13 meta-clones. Clustering features are the time-population-dependent of cell counts for each cell state at each measured time point. This illustrates the distinct terminal fate bias for different meta-clones.

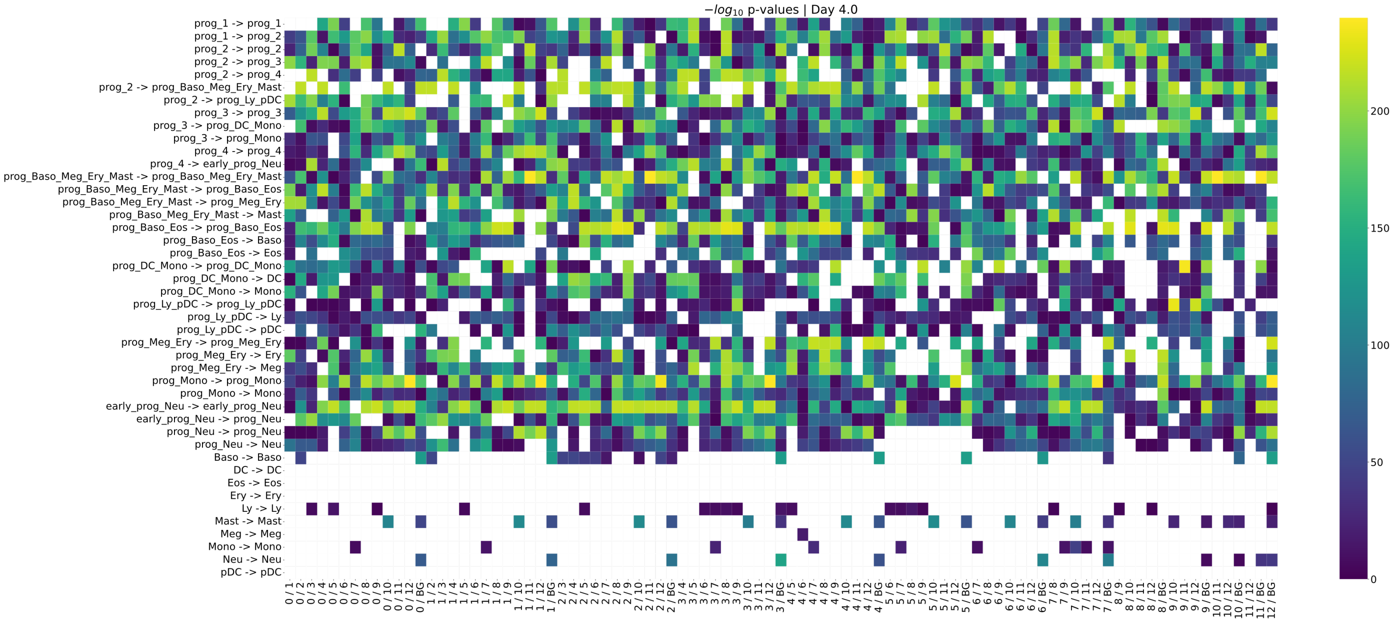


**Figure S13: Statistical comparison of the transition rates for mouse hematopoiesis at Day 4.0.** The student t-test and Mann-Whitney U rank test were used to determine whether the bootstrapping derived rate distributions are significantly different from each pair of meta-clones. We also used the mean absolute change of all bootstrapping trials to further filter outliers. The heatmap uses $-log_{10}(p)$ value for better visualization purpose, therefore, lighter entries represent higher differences between a pair of meta-clones. Blank indicates no significance or the result.

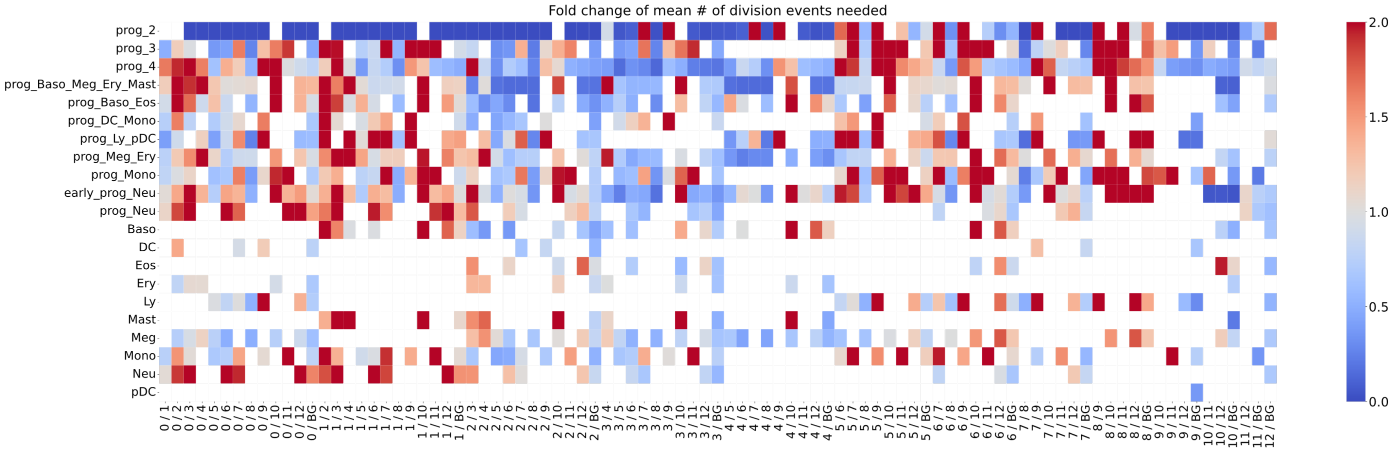


**Figure S14: Fold change comparison of mean number of division events needed to produce the first progeny given a prog_1 for mouse hematopoietic dataset.** Warmer colors indicate a larger number of divisions and the other way round. Blank parts represent no significance or the corresponding meta-clone does not produce the associated populations.

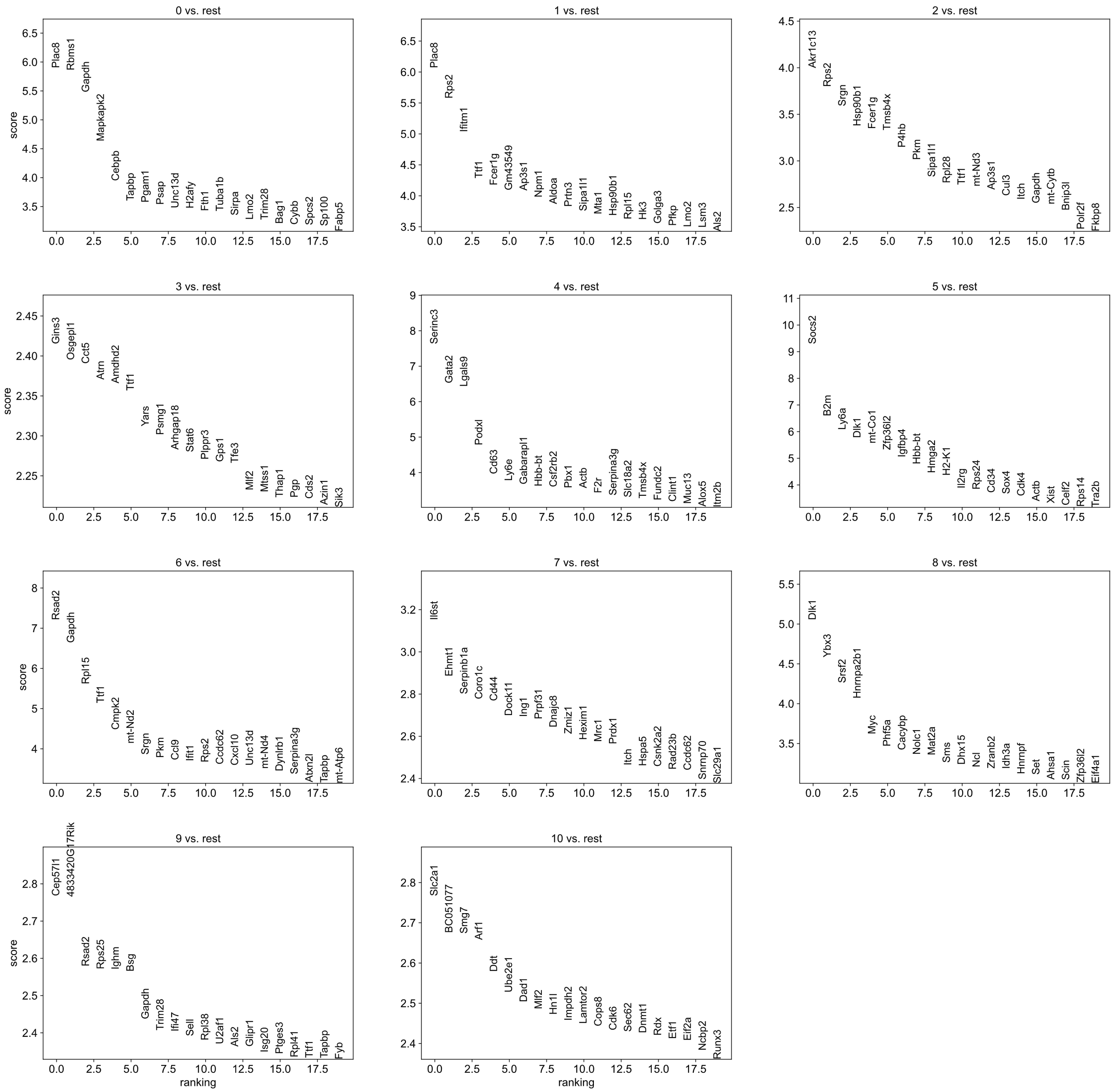

**Figure S15: Differential Expression Genes analysis between all meta-clones using only prog_1 populations of mouse hematopoietis system.** The preference of different lineages of each meta-clone could be revealed in the very early stage of development. Although they were all stem cells, they still have subtle difference in DEGs among each other which indicates their differentiation potential, e.g., multi-potent meta-clones or uni-potent meta-clones.

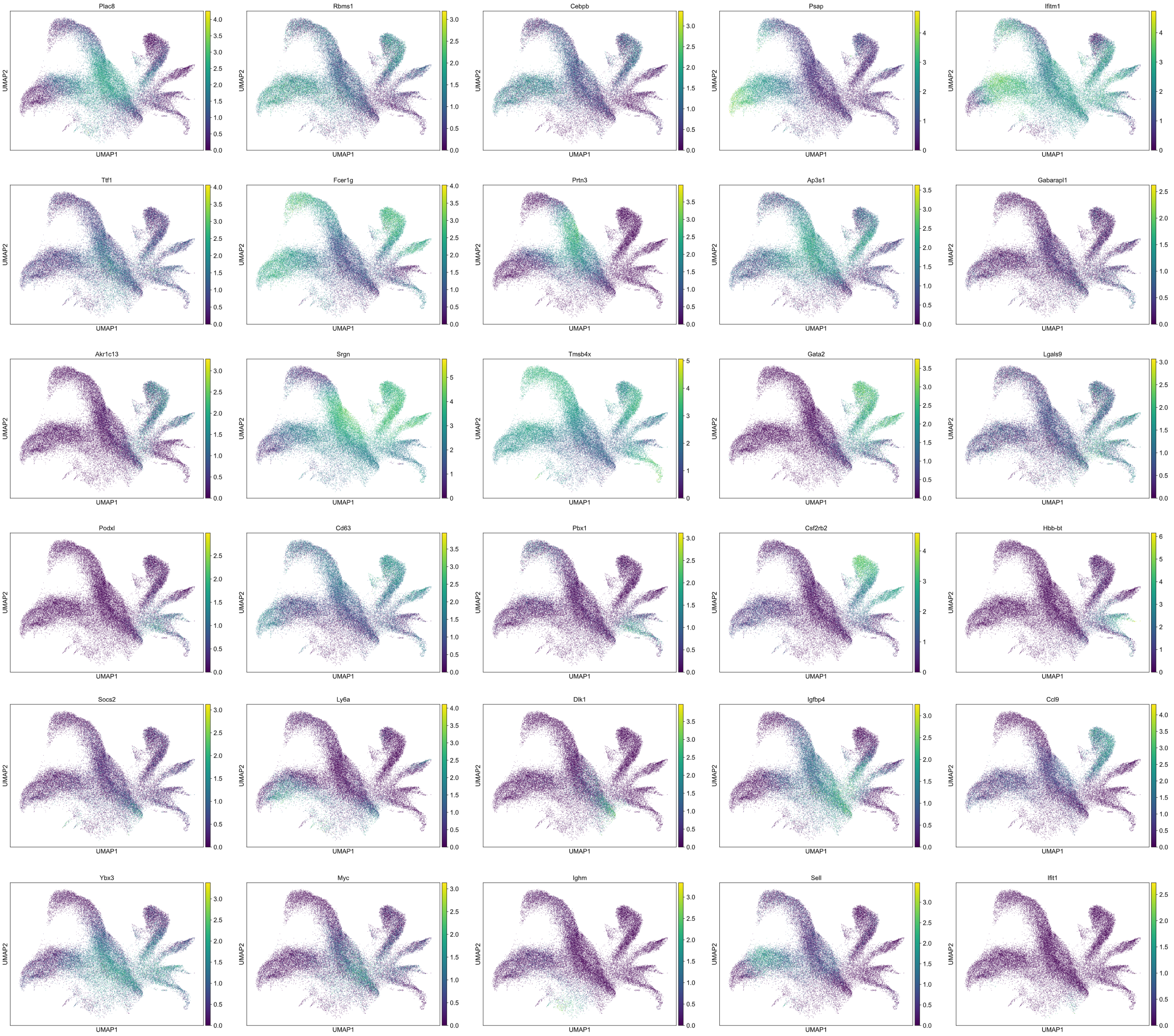


**Figure S16: Selected DEGs exhibits fate bias of progenitor cells and serve as markers for specific lineages.** Note that the DEGs presented here are extracted from Supplementary Fig. S15, which means they the are DEGs between progenitor cells, while their expression is plotted for all cells to highlight lineage preferences.

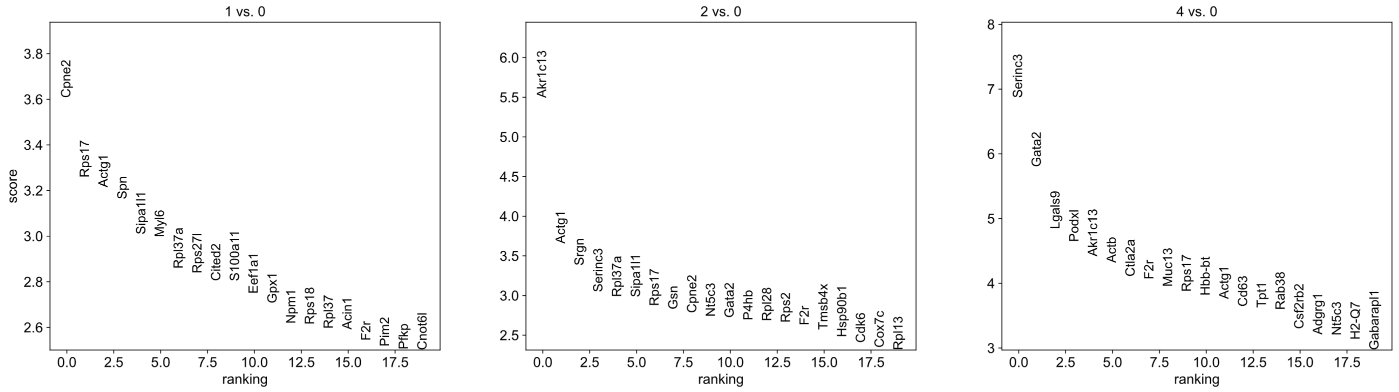


**Figure S17: DEGs analysis between meta-cloens 0~4 using only prog_1 populations of mouse hematopoietis system, with meta-clone 0 as reference population.**

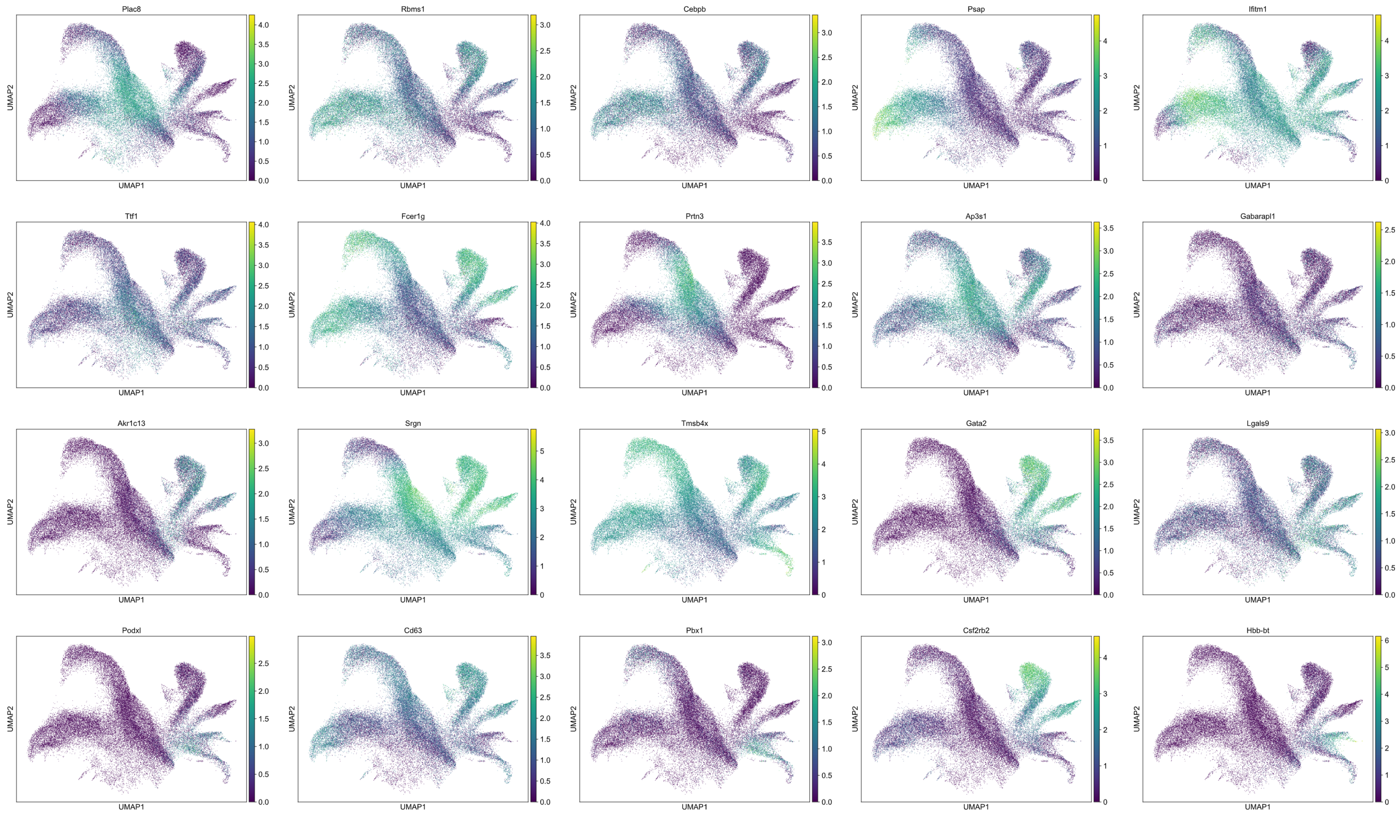


**Figure S18: Selected DEGs exhibits fate bias of progenitor cells and serve as markers for differentiation ability.** The DEGs presented here are extracted from Supplementary Fig. S17, which means they the are DEGs between progenitor cells, while their expression is plotted for all cells to highlight lineage preferences.
