## Supplementary Notes for "Unveiling Clonal Cell Fate and Differentiation Dynamics: A Hybrid NeuralODE-Gillespie Approach"

**Supplementary Note 1: Interpretation of the transition rates between populations.**

Based on expert curated PAGA graph and penalty term as constraints, the per capita and per day change of a specific population is the combined rates (within a row) in the transition matrix. Whilst the model supports two modes, constant mode (time-invariant transition rates) and dynamic mode (time-variant transition rates), we use constant rate as an example for simplicity.

Given a cell population $p_i$, the change in total counts comes from three different aspects: differentiating cells from other populations (inbound), net growth of $p_i$ and differentiation towards other populations (outbound), where the latter two can be combined as total cell loss when considering the overall effect on population size. Therefore, for population with no ancestors such as HSCs, we have an exponential change,

$$N(t) = N(0)\, e^{rt} \tag{1}$$

where $r$ is the per capita net growth rate and $N$ is the population size. On the other hand, for intermediate or terminal cell states, the population size at time $t$ is also affected by other ancestor states as shown:

$$N_i'(t) = \sum_j N_j(t) * K_{ji} - N_i(t) * \sum_j K_{ij} \tag{2}$$

where $i$ and $j$ here are indexes of two populations, hence the meaning of $K$ here is not the per capita growth rate $r$. To illustrate this concept, suppose HSC has 2 progenies (MEMP and Neutrophil-myeloid progenitor) and MEMP has 3 progenies (Megakaryocyte, Mast cell and Early Erythroid). Then the change of population size in HSCs per unit of time is the newly generated HSCs through proliferation process minus number of HSCs turn into MEMP and NMP through differentiation process. Likewise, the change of popualtion size in MEMP per unit of time is calculated based on the similar formulas described above (proliferation / differentiation to other states), in addition to that intermediate cell states (e.g., MEMP) are partially produced by its ancestors (e.g., $N_{hsc} * 2.3$).

$$\begin{aligned} N'_{hsc}(t) &= N_{hsc} * (1.6 - 2.3 - 0.25) \\ &= N_{hsc} * (-0.95) \\ N'_{memp}(t) &= N_{memp} * (2.6 - 0.44 - 0.89 - 1.2) + N_{hsc} * 2.3 \\ &= N_{memp} * 0.07 + N_{hsc} * 2.3 \end{aligned} \tag{3}$$

**Supplementary Note 2: Gillespie algorithm for stochastic simulation of cell divisions.**

A continuous time Markov process can be completely defined by its transition rates between states and their initial conditions. The first moments of the model can be modeled with ordinary differential equations, then the conditional probability and definite state trajectories can be calculated. However, this only gives us a deterministic result of the model with time evolution as a continuous and predictable process. On the contrary, the stochastic approach sees trasitions as a random-walk process; it can be modeled by master equations and yields a more precise and comprehensive model. The master equation represents the evolution process of a multi-state system along time, which can be expressed using the following form:

$$P_t' = K(t)P_t \tag{4}$$

where the transition rate matrix $K(t)$ can either be time-invariant or time-variant and $P_t$ is the probability distribution of states at time $t$.

The Gillespie algorithm (or the Kinetic Monte-Carlo) is a simulation algorithm used to explore the probability space of a Markov process with a given transition matrix; the algorithm allows us to model the dynamics described by the master equations. However, because of the random nature of a stochastic process, the algorithm must be run multiple times so that it produces a set of stochastic trajectories based on random seeds. Within each iteration, a single chain of random events along the time course is created. Theoretically, with a large list of chains combined, the overall states of the system can be captured and the mean is exactly the same with the ODE based model whilst Gillespie provides more information and heterogeneity of the system.

We denote the rate of proliferation and differentiation between cell populations to be $K_{ij}$ and the cell counts as $N_i$. Therefore, the parameter $\lambda$ is calculated by,

$$\lambda = \sum_i \sum_j N_i \cdot k_{ij} \tag{5}$$

which is also called the propensity score and the probability of a given reaction is derived as,

$$P_{ij} = \frac{N_i \cdot k_{ij}}{\lambda} \tag{6}$$

The time until the occurrence of the next event is independent of previously occurred events. Thus, a Poisson process is usually used to satisfy this constraint where the number of events happened during a time interval $\Delta t$ follows a Poisson distribution and subsequently the time interval between two reactions then follows an exponential distribution with

$$\Delta t = \frac{ln(1/u_0)}{\lambda} \tag{7}$$

where $u_0$ is a random number sampled from $u \in U(0,1)$. The complete pseudo-code of Gillespie algorithm for modeling cell division is shown in algorithm 1.

**Algorithm 1:** Gillespie algorithm for stochastic simulation of cell divisions

**Data:** Time $t$, cell counts $N_i$, transition matrix $K_{ij}$, total number of cells produced $I_i$, $i$: the index of states
**Result:** Stochastic description of a biological process, number of divisions between a HSC and a progeny

**for** $iter \leftarrow 0$ **to** $1,000$ **do**
  Initial $N_{hsc} \leftarrow 1, N_{others} \leftarrow 0, I_i \leftarrow 0, t \leftarrow 0$;
  **while** $t \leq t_{max}$ *and* $\sum_i N_i > 0$ **do**
    Compute the total propensity score $\lambda = \sum_i \sum_j N_i \cdot k_{ij}$;
    Generate uniformly distributed numbers $\{u_0, u_1\}$ randomly from $u \in U(0,1)$;
    Calculate time increment of when the next reaction will occur $\Delta t = \frac{ln(1/u_0)}{\lambda}$;
    Select the reaction $q$ using $u_1$ with weighted probability $p_{ij} = \frac{N_i \cdot k_{ij}}{\lambda}$ ;
    Pick a cell $n_i$ randomly from $N_i$ that is involved in reaction $q$;
    Update list of cell counts $N_i$ and maximum ID $I_i$ according to the outcome of reaction $q$;
    **Delete** $n_i$;
    $t \leftarrow t + \Delta t$;
  **end**
**end**

**Supplementary Note 3: Laboratory methods for the generation of LARRY-barcoded cord blood progenitor cell datasets.**

**1) LARRY barcoding plasmid propagation and analysis of barcode library diversity using 10X.**

The LARRY Barcode Version 1 library was a gift from Fernando Camargo (Addgene, #140024). The library was amplified and the lentiviral vector produced according to the associated published protocol[18] with some minor adaptations. Briefly, plasmids were introduced into ElectroMAX Stbl4 competent cells (Life Technologies) using MicroPulser Electroporator (Biorad) and incubated for 1 hour at $37°C$ before spreading over 24 large Agar + Ampicillin plates. After 24 hours at $32°C$, colonies were harvested through scraping using pre-warmed LB medium containing Ampicillin. The resulting culture (approx. 1.5L) was incubated at $37°C$ for 2 hours before isolating plasmid DNA using Megaprep kits (Machery-Nagel).

A reference library was made through sequencing of PCR-amplified barcodes from the LARRY plasmid library. $10ng$ of LARRY plasmids was taken as input for a two-step PCR; the first step adds Illumina Read1 and Read2 sequences (5'ACACTCTTTCCCTACACG ACGCTCTTCCGATCTTGTGACGTCACAGGTCGACACCAGTCTCATT3' and 5'GTGACTGGAGTTCAGACGTGTGCTCTTCCGATCGAGTAACCGTTGCTAGGAGAGACCATA3'). The second step adds the P5 and P7 flow cell attachment sequences and 7bp sample indices (P5 5'AATGATACGGCGACCACCGAGATCTACACTCTTTCCCTACACGACGCTCTTCCGATCT3' and P7 5'CAAGCAGAAGACGGCATACGAGANNNNNNNGTGACTGGAGTTCAGACGTGCTCTTCCGATC3'). $8ng$ of PCR1 product was taken as input for PCR2. The PCR programme used was as follows: $98°C$ 2 min, 8 cycles of $98°C$ 10 sec, $58°C$ (PCR1), $62.5°C$ (PCR2) 20 sec, $72°C$ 30 sec, followed by final elongation $72°C$ 5 min and $4°C$ indefinitely. In between PCR1 and PCR2, PCR purification was performed using the QIAquick PCR purification kit (Qiagen). Purification of PCR2 product was carried out using Ampure XP beads (Beckman Coulter) before sequencing on a Novaseq instrument at the Cancer Research UK Cambridge Institute Genomics facility. The resulting list of LARRY barcodes were used as a reference list in downstream analysis.

**2) LARRY library lentivirus preparation.**

Lentiviral vector was produced by transforming the amplified LARRY and packaging plasmids (psPAX2 and pMD2.G) into HEK293T cells using the TransIT-LT1 transfection reagent (Mirus) and incubated at $37°C$. 24 hours after transfection, $500\mu l$ of 0.5 mM Sodium Butyrate (Sigma-Aldrich) was added to the cells and the culture continued at $37°C$ incubation. LARRY lentivirus was harvested 48 hours after transfection, filtered through a $0.45\mu m$ PES filter (Whatman) and concentrated 100-fold by centrifugation in a Beckman Coulter Optima XPM-80 ultracentrifuge at 20,000 x g for 2 hours at $4°C$. The lentivirus was aliquoted and stored at $-80°C$. Titration of the lentiviral barcoding library was performed on HEK293T cells, with a read out obtained by flow cytometry 3 days after infection.

**3) Cord blood samples.**

Umbilical cord blood (CB) samples were obtained with informed consent from healthy donors by Cambridge Blood and Stem Cell Biobank (CBSB) in accordance with regulated procedures approved by the relevant Research and Ethics Committees (18/EE/0199 Research Study). MNCs were obtained using Pancoll density gradient centrifugation of diluted (1:1 with PBS) CB. Red cells were lysed before positive selection for CD34$^+$ cells using the micro beads CD34$^+$ selection kit and AutoMACS cell separation technology (Miltenyi biotech). CB CD34$^+$ cells were then stored at $-150°C$ until use in experiments.

**4) FACS.**

To sort CD34$^+$ cells for experiments from CB samples, the cells were thawed by dropwise addition of pre-warmed Rich Thawing Medium containing Iscove's Modified Dulbecco's Medium (IMDM, Life Technologies), 0.1mg/ml DNase (Lorne Laboratories) and 50% Fetal Bovine Serum (FBS, Life Technologies) before resuspending in PBS + 3% FBS. The samples were stained with the following antibodies:

| | | | | |
|---|---|---|---|---|
| Marker | Lineage | CD11c | CD38 | CD34 |
| Fluorochrome | FITC | FITC | PECy7 | APC |
| Dilution | 1 in 100 (Biolegend) | 1 in 100 (Biolegend) | 1 in 100 (Biolegend) | 1 in 200 (BD) |

Cells were incubated for 20 minutes at room temperature and washed with PBS + 3% FBS. DAPI (Biolegend) was added 1 in 100 (final concentration $137\mu g/ml$) to identify viable cells. Unstained cells and compensation beads (Invitrogen) were used for compensation and as controls to set appropriate gates. Lineage$^-$ CD11c$^-$ CD34$^+$ cells were sorted into Eppendorf tubes using a BD Influx approved for CL2-sorting.

**5) In vitro barcoding, culture and sampling for 10X.**

CB CD34$^+$ cells were cultured in previously published conditions[74], that promote the differentiation of cells towards Megakaryocyte, Erythroid and Myeloid lineages (MEM) with reduced levels of EPO cytokine to avoid biasing the culture towards the erythroid lineage.

Reduced EPO MEM culture media recipe:

| **Stem Pro** | **(mL)** | **Ratio to media (per mL media)** | **Final (mL)** |
|---|---|---|---|
| Media | 40 | | 50 |
| Nutrients | 1.4 | 0.035 | 1.75 |
| L-Glutamine | 0.5 | 0.01 | 0.5 |
| P/S | 0.5 | 0.01 | 0.5 |

| | **Final (ng/mL)** | **Stock (ng/$\mu$L)** | **Volume ($\mu$L)** |
|---|---|---|---|
| SCF | 100 | 100 | 50 |
| Flt3 | 20 | 50 | 20 |
| TPO | 100 | 50 | 100 |
| EPO | 2 units | 2 | 50 |
| IL-6 | 50 | 50 | 50 |
| IL-3 | 10 | 50 | 10 |
| IL-11 | 50 | 50 | 50 |
| GM-CSF | 20 | 50 | 20 |
| IL-2 | 10 | 50 | 10 |
| IL-7 | 20 | 40 | 25 |
| Lipids | 50 | 50 | 50 |

For barcode labelling, 75,000 sorted Lineage$^-$ CD11c$^-$ CD34$^+$ cells were seeded at 5,000 cells per well in a 96 well round-bottom plate. LARRY lentivirus was added directly to the culture at an MOI of 60. After 24 hours, the virus was diluted out and cells transferred to a 96 well flat-bottom plate. At 3 days post-transduction, GFP$^+$ and GFP$^-$ cells were FACS-sorted. Two thirds of the GFP$^+$ fraction (8,856 cells) was sent for scRNA-seq analysis (10X genomics) and the remaining third (4,428 cells) was re-plated in 500 $\mu l$ reduced EPO MEM culture conditions in a 24 well plate. GFP$^-$ cells were also analysed by scRNA-seq. At day 10 post-transduction cells were stained for GlyA (PE, 1:1000 dilution, BD) using the protocol above. GFP$^+$ GlyA$^-$ cells were FACS-sorted, with 40,000 cells processed for scRNA-seq and the remainder replated, as previously. At day 17 post-transduction, 40,000 GFP$^{hi}$ GlyA$^-$ and 40,000 GFP$^{mid}$ GlyA$^-$ cells were FACS-sorted and processed for scRNA-seq.

**6) Sample preparation and LARRY cDNA enrichment for scRNAseq (10X Genomics).**

Up to 20,000 live cells of interest were sorted into 300$\mu l$ MEM media and kept on ice before centrifugation. Cells were resuspended in PBS + 0.04% BSA (miltenyi) and further processed for single cell sequencing using the Chromium Single Cell 3' Library & Gel Bead Kit v3 (10X Genomics) following manufacturer's protocols. LARRY molecules constitute a small fraction of the total cDNA library thus we adapted previously published protocols[18,75] to enrich the LARRY barcodes and ensure a sufficient number of reads per barcode. Using a portion of each sample's cDNA, LARRY sequences were PCR-amplified while simultaneously adding Illumina primers and indices necessary for sequencing and identifying the enriched samples in downstream analysis (see table below for primer sequences used). Purification of the PCR product was carried out using Ampure XP beads (Beckman Coulter) before sequencing alongside the full sample cDNA library on a NovaSeq 6,000 at the Cancer Research UK Cambridge Institute Genomics facility.

| Name | Sequence | Sample |
|---|---|---|
| Larry_cDNA_SITTA1_i5 | AATGATACGGCGACCACCGAGATCTACACAGTGTTACCTACACTCTTTCCCTACACGACGCTCTTCCGATCT | Day 3 GFP$^{+}$ |
| Larry_cDNA_SITTA1_i7 | CAAGCAGAAGACGGCATACGAGATCGCATGTTACGTGACTGGAGTTCAGACGTGTGCTCTTCCGATCTACCGTTGCTAGGAGAGACCATATG | Day 3 GFP$^{+}$ |
| Larry_cDNA_SITTA3_i5 | AATGATACGGCGACCACCGAGATCTACACATCAGTCTAAACACTCTTTCCCTACACGACGCTCTTCCGATCT | Day 10 GFP$^{+}$ |
| Larry_cDNA_SITTA3_i7 | CAAGCAGAAGACGGCATACGAGATTTTCGTAGTGGTGACTGGAGTTCAGACGTGTGCTCTTCCGATCTACCGTTGCTAGGAGAGACCATATG | Day 10 GFP$^{+}$ |
| Larry_cDNA_SITTA6_i5 | AATGATACGGCGACCACCGAGATCTACACGAAGTTAGGGACACTCTTTCCCTACACGACGCTCTTCCGATCT | Day 17 GFP$^{high}$ |
| Larry_cDNA_SITTA6_i7 | CAAGCAGAAGACGGCATACGAGATTCACGCGTTAGTGACTGGAGTTCAGACGTGTGCTCTTCCGATCTACCGTTGCTAGGAGAGACCATATG | Day 17 GFP$^{high}$ |
| Larry_cDNA_SITTA7_i5 | AATGATACGGCGACCACCGAGATCTACACAAAGGTAGTAACACTCTTTCCCTACACGACGCTCTTCCGATCT | Day 17 GFP$^{mid}$ |
| Larry_cDNA_SITTA7_i7 | CAAGCAGAAGACGGCATACGAGATACCCTTGGGAGTGACTGGAGTTCAGACGTGTGCTCTTCCGATCTACCGTTGCTAGGAGAGACCATATG | Day 17 GFP$^{mid}$ |
